# Negative modulation of macroautophagy by HERPUD1 is counteracted by an increased ER-lysosomal network with impact in drug-induced stress cell survival

**DOI:** 10.1101/2021.06.14.447273

**Authors:** G. Vargas, O. Cortés, E. Arias-Muñoz, S. Hernández, C. Cerda-Troncoso, L. Hernández, A. E. González, M. H. Tatham, H. Bustamante, C. Retamal, J. Cancino, M. Varas-Godoy, R. T. Hay, A. Rojas-Fernandez, V. A. Cavieres, P. V. Burgos

## Abstract

Macroautophagy and the ubiquitin proteasome system work as an interconnected network in the maintenance of cellular homeostasis. Indeed, efficient activation of macroautophagy upon nutritional deprivation is sustained by degradation of preexisting proteins by the proteasome. However, the specific substrates that are degraded by the proteasome in order to activate macroautophagy are currently unknown. By quantitative proteomic analysis we identified several proteins downregulated in response to starvation but independently of ATG5 expression. Among them, the most significant was HERPUD1, an ER protein of short-half life and a well-known substrate of the proteasome. We found that increased HERPUD1 stability by deletion of its ubiquitin-like domain (UBL) plays a negative role on basal and induced macroautophagy. Moreover, we found it triggers ER expansion by reordering the ER in crystalloid structures, but in the absence of unfolded protein response activation. Surprisingly, we found ER expansion led to an increase in the number and function of lysosomes establishing a tight network with the presence of membrane-contact sites. Importantly, a phosphomimetic S59D mutation within the UBL mimics UBL deletion on its stability and the ER-lysosomal network expansion revealing an increase of cell survival under stress conditions. Altogether, we propose stabilized HERPUD1 downregulates macroautophagy favoring instead a closed interplay between the ER and lysosomes with consequences in drug-cell stress survival.

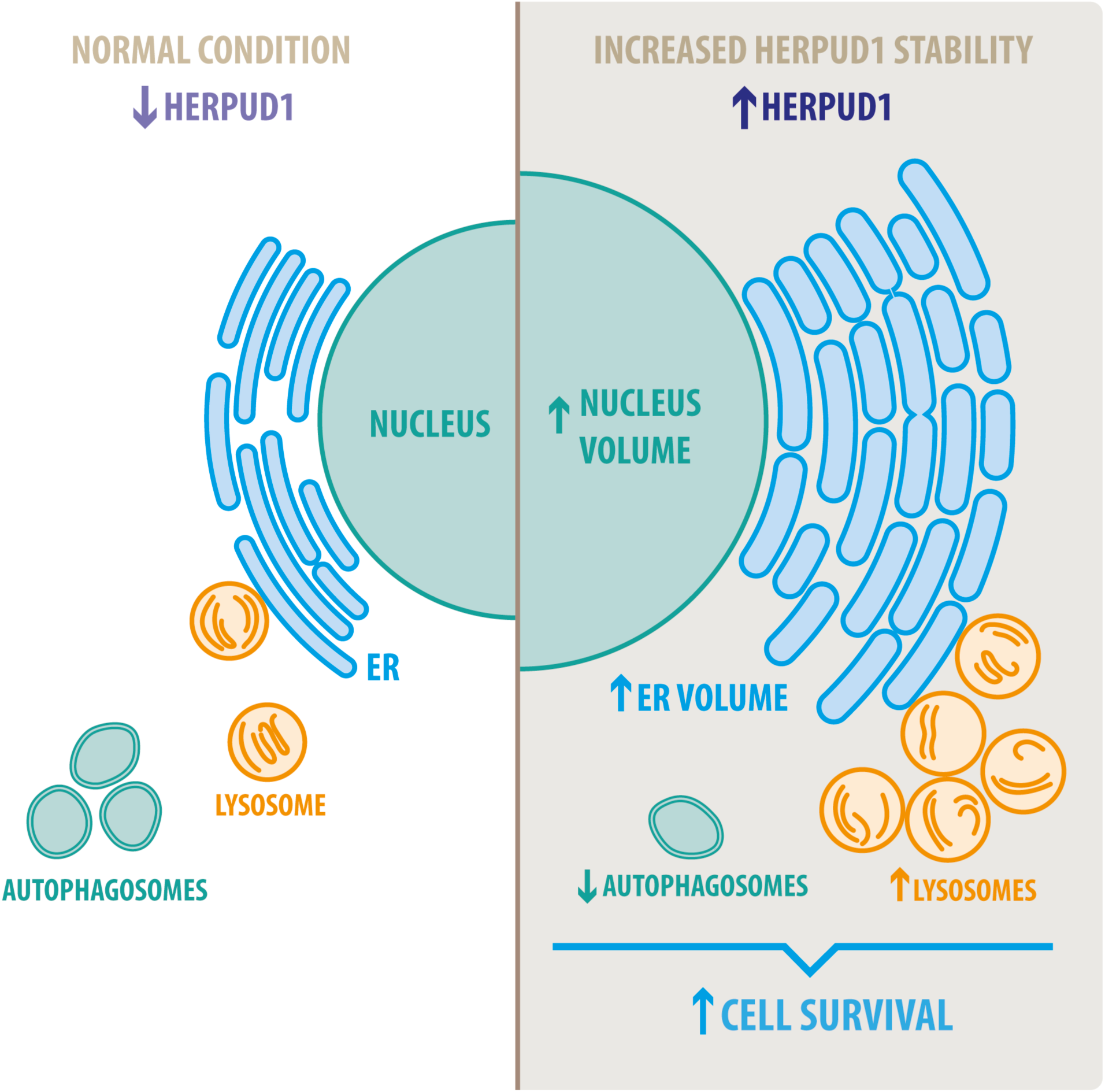

## INTRODUCTION

Macroautophagy (from here referred to as autophagy) is a catabolic pathway that mediates the engulfment of aberrant or damaged cytoplasmic constituents into double-membrane autophagosomes that subsequently fuse with lysosomes to form a hybrid organelle called the autolysosome that mediates the degradation of the cargo by acid hydrolases (Mizushima et al. 2008; Khaminets, Behl, and Dikic 2016). Autophagy is also implicated in the degradation of cellular constituents under basal conditions, playing an essential role in the maintenance of cellular homeostasis upon a variety of environmental conditions such as nutrient restriction or other stressors (Murrow and Debnath 2013). Autophagy is highly inducible by environmental changes being a highly dynamic process that resolves a variety of cellular demands (Murrow and Debnath 2013). In fact, increased autophagy is protective in different cells and organisms, playing a crucial role in cell maintenance and survival under different insults (Moreau, Luo, and Rubinsztein 2010). On the other hand, defects in autophagy enhance cell vulnerability under harmful conditions such as those present in the tumor microenvironment (Camuzard et al. 2020).

Although initially autophagy was thought to work independently of the ubiquitin proteasome system (UPS), increasing evidence shows many layers of both negative and positive regulation (Bustamante et al. 2018), revealing an interconnected network with important roles in cellular homeostasis and maintenance (Korolchuk, Menzies, and Rubinsztein 2010). Inhibitors of the proteasome 20S catalytic core with the use of β-subunits blockers triggers an enhancement in the biogenesis of autophagosomes (Zhu, Dunner, and McConkey 2010). In contrast, impairment of the proteasome 19S regulatory particle with an inhibitor of PSMD14, a proteasomal deubiquitinating enzyme, blocks the biogenesis of autophagosomes (Demishtein et al. 2017; Bustamante et al. 2020). To date a limited number of substrates of the UPS system are known to play a regulatory role in autophagy (Jia and Bonifacino 2019; Thayer et al. 2020) and many aspects about the functional role of this interconnected network between autophagy and UPS remain elusive.

To identify UPS substrates that could act as negative regulators of autophagy, we conducted a quantitative SILAC proteomic analysis in cells stably depleted of ATG5 by shRNA-mediated knockdown. ATG5 protein is part of a complex with ATG12 and ATG16L that controls an essential step in the autophagosome formation (Walczak and Martens 2013). We focused on proteins downregulated in response to nutrient deprivation, but not because of autophagy activation. The protein with the most significant downregulation, in wild type and ATG5 depleted cells was the Homocysteine-responsive endoplasmic reticulum-resident ubiquitin-like domain (UBL) member 1 protein named as HERPUD1. This protein is a transmembrane ER-resident protein with low levels of expression due to its short half-life by rapid proteasomal degradation (K. Kokame et al. 2000; Sai et al. 2003).

Here, we found that increased HERPUD1 stability through the deletion of its UBL domain causes a decrease in basal and induced autophagy. Additionally, it promotes an ER expansion which is organized in stacked tubules forming crystalloid ER-like structures but independent of ER stress. Furthermore, we uncovered that higher HERPUD1 stability has a positive impact in lysosomal function, establishing an ER-lysosomal network with the presence of membrane contact-sites (MCS). Further, combining bioinformatics and site-directed mutagenesis we found the phosphomimetic S59D mutant within the UBL domain of HERPUD1 mimics the effect of the UBL deletion increasing its levels and inducing the expansion of the ER-lysosomal network, a phenotype that promotes stress cell survival. These findings thus identify HERPUD1 as a hotspot platform to promote stress cell survival by inducing an expansion of the ER-lysosomal network when autophagy slows down.

## MATERIALS AND METHODS

### Reagents

Bafilomycin A1 (BafA1, cat#B1793), tunicamycin (Tun, cat#T7765), thapsigargin (Tg, cat#T9033), cisplatin (CDDP, cat#479306), Sulforhodamine B (SRB, cat#230162), Earle’s balanced salt solution (EBSS, Cat#E2888), puromycin dihydro-chloride (cat#P8833), and protease inhibitors cocktail (cat#P8340) were purchased from Sigma-Aldrich (St. Louis, MO, USA). MG132 (cat#474790) was purchased from Merck Millipore (Burlington, MA, USA). LysoTracker^TM^ Red DND-99 (cat#L7528), 4′,6-diamidino-2-phenylindole (DAPI) (cat#D-1306) and TRIzol^TM^ (cat#15596018) were purchased from ThermoFisher Scientific (Waltham, MA, USA), Magic Red® (cat#6133) was purchased from Immunochemistry Technologies, LLC (Bloomington, IN, USA). The QuikChange II XL direct-mutagenesis kit was obtained from Stratagene (cat#200522, La Jolla, CA, USA) and the Vybrant Apoptosis Pacific Blue-annexin V kit and 7AAD from Invitrogen (cat#A35122).

### Antibodies

The following monoclonal antibodies were used: mouse anti-β-ACTIN clone BA3R (cat# MA5-15739, Thermo Fisher Scientific), mouse anti-XBP-1S clone E7M5C (cat#47134S, Cell Signaling Technology, Danvers, MA, USA), mouse anti-FLAG clone M2 (cat#F1804, Sigma Aldrich), mouse anti-GRP78/BiP clone 40/BiP (cat# 610978, BD Biosciences, San Jose, CA, USA), mouse anti-LAMP1 clone H4A3 (cat# 610978, Developmental Studies Hybridoma Bank, Iowa City, IA, USA), rabbit monoclonal anti-CALNEXIN clone C5C9 (cat#2679S, Cell Signaling Technologies), rabbit monoclonal anti-ATF4 clone D4B8 (cat#11815S, Cell Signaling Technologies), rabbit monoclonal anti-HERPUD1 clone EPR9649 (cat#ab150424, Abcam), rat monoclonal anti-GRP94 clone SPM249 (cat#ab233979, Abcam, Cambridge, UK). We used the following polyclonal antibodies: rabbit anti-LC3 (cat#2775S, Cell Signaling Technology), rabbit anti-PERK (cat#P0074, Sigma-Aldrich), goat anti-CATHEPSIN-D (cat#AF1014, R&D Systems, Minneapolis, MN, USA), rabbit anti-HERPUD1 (cat#BML-PW9705, ENZO Life Sciences, Farmingdale, NY, USA). Horseradish peroxidase-conjugated secondary antibodies were purchased from Jackson ImmunoResearch Laboratories (West Grove, PA, USA), Alexa fluorophore-conjugated secondary antibodies were purchased from Thermo Fisher Scientific.

### Mass Spectrometry based proteomics and statistical analysis

Human H4 neuroglioma cells stably expressing shLuc and shATG5 were grown in Dulbecco’s modified Eagle’s medium lacking all amino acids except L-lysine and L-arginine, which were replaced with either unlabelled amino-acids (Lys0 and Arg0) or stable isotopic forms 13C6 15N2-lysine (Lys8) and 13C6 15N4-arginine (Arg10) (Cambridge Isotope Laboratories). The medium was supplemented with 10% dialyzed FBS using previous published methods (Golebiowski et al. 2009; Yin et al. 2012). Two separate cultures of each shLuc and shATG5 cells were grown in light amino acids (Lys0 + Arg0) or heavy amino acids (Lys8 + Arg10) over 9 replication cycles to achieve over 96% incorporation and similar cell counts per culture. Both cell cultures were divided in two, and cells were grown under control conditions in DMEM + 10% FBS or starvation in EBSS media for 4 h so that each cell type grown with each label were treated with both conditions (8 cultures in total). Cells were washed and lysed in 1.2 x LDS sample buffer with reducing agent, obtaining crude cell extracts at approximately 1mg/ml. To allow all experimental conditions to be compared with one another a light reference mix was obtained by combining all four light amino acid conditions in a 1:1:1:1 ratio by volume. This was then mixed 1:1 ratio (v:v) with each heavy amino acid extract. The same comparisons were made in reverse by combining all heavy amino acid samples into a reference mix and combining this 1:1 (v:v) with each of the four individual light amino acid conditions. See Figure 1A for experimental design. All 8 mixes were fractionated by SDS-PAGE and stained gels were cut into 3 slices per lane before tryptic peptides were extracted. The resultant 24 samples of dried down peptides were resuspended in 35 µl 0.1% TFA 0.5% acetic acid. Peptide samples were analyzed by LC-MS/MS twice; the first using 9 µl peptide sample run over a 90 min peptide fractionation gradient, and the second using 18 µl peptide sample run over a 240 min fractionation gradient. Peptides were analyzed using a Q exactive mass spectrometer (Thermo Scientific) coupled to an EASY-nLC 1000 liquid chromatography system (Thermo Scientific), using an EASY-Spray ion source (Thermo Scientific), running a 75 µm x 500 mm EASY-Spray column at 45°C. A top 10 data-dependent method was applied. Full scan spectra (m/z 300-1800) were acquired with resolution R= 70,000 at m/z 200 (after accumulation to a target value of 1,000,000 with maximum injection time of 20ms). The most intense ions were fragmented by HCD and measured with a resolution of R= 17,500 at m/z 200 (target value of 500,000 maximum injection time of 60 ms) and intensity threshold of 2.1×10^4^. Peptide match was set to “preferred”, a 40 second dynamic exclusion list was applied, and ions were ignored if they had an assigned charge state of 1, 8 or >8. All 48 data files were analyzed simultaneously in MaxQuant (v1.5.2.8) using default parameters excepting the selection of SILAC labels, activation of ‘requantify’ and ‘match between runs’, using a uniport ‘HUMAN’ proteome database (downloaded 24/02/2015- 73920 entries) as search space. Raw files derived from the same mix were grouped under the ‘Experiment’ heading as Mix01-Mix08. Raw files were given MaxQuant experimental design ‘fraction’ numbers such that spectra derived from the same HPLC gradient and from the equivalent gel slices across different lanes would be matched, as well as one sliced either side. 4395 protein groups were listed as informed in the Supp. File 1, proteinGroups.txt sheet. This list was edited to leave 3911 protein groups by removing decoy proteins, protein listed as potential contaminants, proteins identified only by modified peptides, and proteins without a H/L ratio reported for any comparison (see Supp. File 1, accepted sheet). The average of normalized forward and reverse ratios of starvation/control were carried forward for statistical analysis, but only for those with H/L ratios reported for both. These values along with average log_10_ protein intensities were used in Perseus to calculate Significance B values to identify statistical outliers (SigB<0.001) for the three ratios. Data for these shortlisted proteins are summarized in Supp. File 1, shortlisted sheet. Data is available via ProteomeXchange with identifier PXD024486. Reviewer account details: Username: reviewer Password: ibjQUyjo

**Figure 1.**
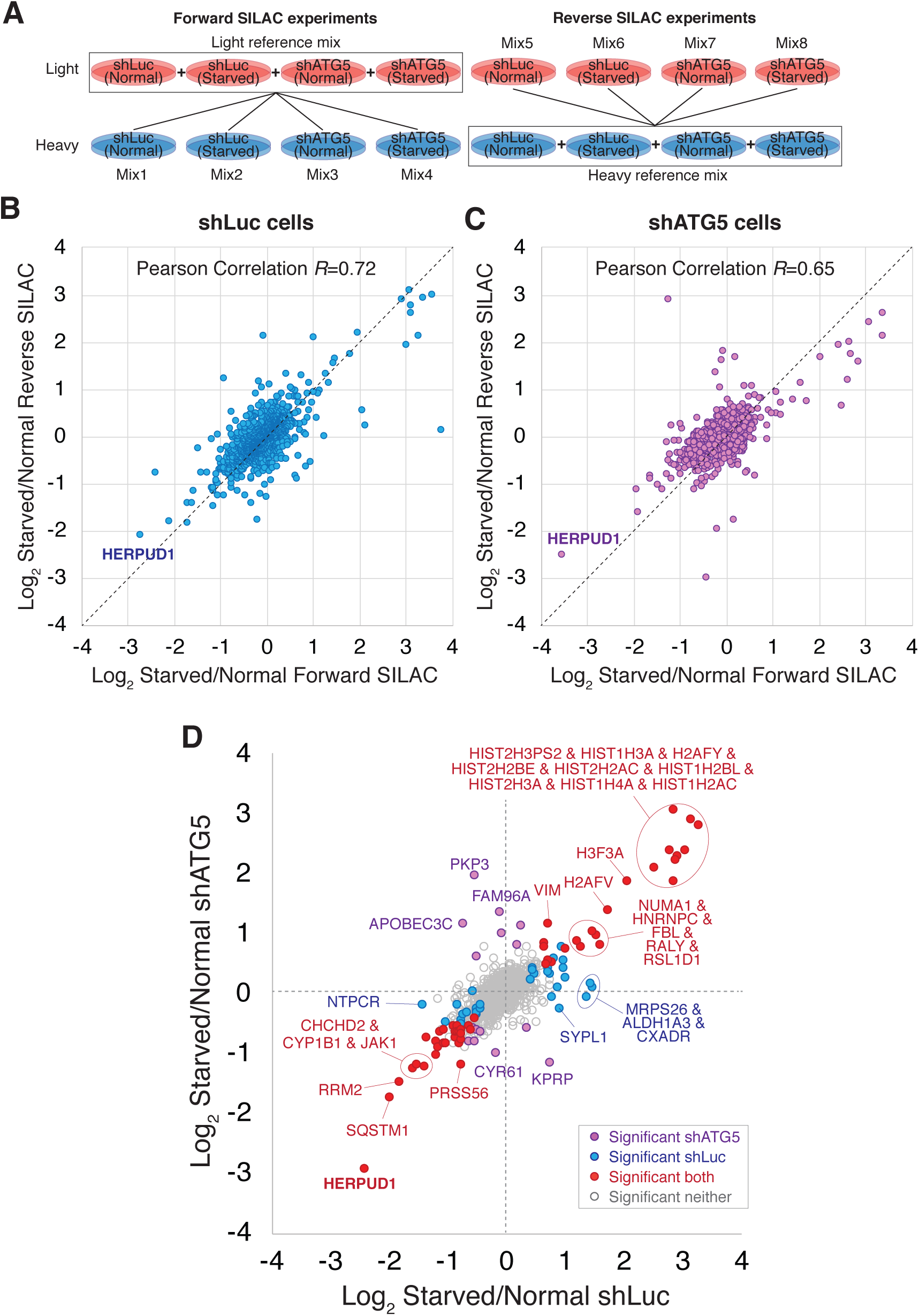
SILAC-based proteomic study reveals HERPUD1 as a possible modulator of autophagy. **(A)**. Design of the SILAC experiment to monitor changes of the cellular proteome of H4 cells during starvation and shRNA-mediated knockdown of ATG5 (shATG5) or shRNA against the luciferase gene (shLuc). ‘Reference’ mixes of all cell extracts were prepared for light and heavy conditions for ‘forward’ and ‘reverse’ SILAC experiments and were mixed 1:1 with individual cell extracts, giving a total of 8 mixes. **(B-C)** Scatter plot comparing ‘Forward’ and ‘Reverse’ Log2 Starvation/Normal ratio data for cells shLuc (B) and shATG5 (C). **(D)** Comparison of average Log_2_ Starvation/Normal ratios in shLuc (x-axis) and shATG5 (y-axis). Protein outliers under either or both knockdown conditions are colored as indicated.

### Plasmids and site-directed mutagenesis

For all HERPUD1 constructs generated in this study, previously described cDNAs encoding either the full-length or the ΔUBL deletion mutant human HERPUD1, both with a C-terminal FLAG-tag, were used as templates (Sai et al. 2003). pCI-HERPUD1-FLAG and pCI-ΔUBL-FLAG were digested with EcoRI and NotI restriction enzymes to obtain an insert, which was sub-cloned to a lentiviral pLVX-IRES-Puro vector (Takara Bio Inc, CA, USA) containing a puromycin resistance gene. The substitution S59D and S59A were introduced into the pLVX-IRES-FLAG-tagged HERPUD1 vector using the QuikChange II XL direct-mutagenesis kit (Stratagene, cat#200522) and the mutagenesis service of GenScript (Hong Kong, China).

### Cell culture and generation of stable cell lines

Maintenance of H4 human neuroglioma cells stably expressing either shRNAs against ATG5 or luciferase genes was performed as previously described (González et al. 2017). HeLa cells were obtained from the American Type Culture Collection (Manassas, VA, USA). HeLa-derived cell lines were cultured in Dulbecco’s modified Eagle’s medium (DMEM; Thermo Fisher Scientific) supplemented with 10% (vol/vol) heat-inactivated fetal bovine serum (FBS; Thermo Fisher Scientific), and 100 U/ml penicillin/100 mg/ml streptomycin (Thermo Fisher Scientific), in a 5% CO_2_ atmosphere at 37°C. The generation of HeLa stable cell lines expressing all different variants of FLAG-tagged HERPUD1 cloned in the pLVX-IRES-Puro vector were generated by transfection with Lipofectamine 2000 (Invitrogen) according to manufacturer’s instructions. After 24 h the cells were selected and maintained with 2 µg/ml of puromycin.

### Preparation of protein extracts, Electrophoresis, SDS-PAGE and Western blot analysis

Cells were washed with ice-cold phosphate buffered saline (PBS) and lysed in Radioimmunoprecipitation assay buffer (RIPA lysis buffer) [50 mM Tris-HCl, 150 mM NaCl, 5 mM EDTA, 1% NP-40, 1% sodium deoxycholate, 0.1% SDS, pH 7.4], supplemented with a cocktail of protease inhibitors [416 µM 4-(2-Aminoethyl)benzenesulfonyl fluoride, 0.32 µM Aprotinin, 16 µM Bestatin, 5.6 µM E-64, 8 µM Leupeptin and 6 µM Pepstatin A; Sigma-Aldrich] and phosphatase inhibitors (1 mM NaF, 0,3 mM Na_2_P_2_O_7_ and 1 mM Na_3_VO_4_; Sigma-Aldrich). Cell lysates were collected and lysed for 30 min at 4°C in rotation. Then, extracts were sonicated with ultrasonic power three times on ice with pulses of 2-3 sec at 40 mA using a tip sonicator system. Extracts were further centrifuged for 20 min at 13.000xg at 4°C. The supernatants were collected, and protein concentration was quantified using the BCA assay (ThermoFisher Scientific). The protein extracts were denatured at 65°C for 5 min and analyzed using our previous described methods (González et al. 2017; Bustamante et al. 2020).

### Transmission electron microscopy

HeLa cells were fixed for 16 h by immersion in 2.5% glutaraldehyde in 0.1 M cacodylate buffer (pH 7.0) at room temperature, and then washed three times with a cacodylate buffer for 2 h. Cells were post-fixed with 1% osmium tetroxide (OsO_4_) for 2 h and washed 3 times with double distilled water. Then, the cells were treated with 1% aqueous uranyl for 90 min, and sequentially dehydrated through an acetone battery 50%, 70%, 95%, and 100% for 20 min each. Cells were pre-embedded in epon/acetone 1:1 overnight and then in pure epon for 4 h. Finally, cells were embedded in fresh resin and polymerized in an oven at 60°C for 48 h. Ultrafine sections (80 nm) were obtained using an ultramicrotome Leica Ultracut R. The sections were incubated with 4% uranyl acetate in methanol for 2 min and lead citrate for 5 min. The grids were visualized using a Philips Tecnai 12 electron microscope (Eindhoven, The Netherlands) at 80 kV. This work was performed in the Advanced Microscopy Facility of the Faculty of Biological Sciences at Pontificia Universidad Católica de Chile.

### Fluorescence microscopy, data acquisition and quantification

Cells grown on glass coverslips were washed with PBS-Ca^2+^/Mg^2+^ and then fixed, permeabilized and stained using our published protocols (Bustamante et al. 2020; Cavieres et al. 2020). The images were acquired using a TCS SP8 laser-scanning confocal microscope (Leica Microsystems, Wetzlar, Germany) equipped with a 63× oil immersion objective (1.4 NA), photomultipliers (PMT), hybrid detectors (HyD) system using 405 nm, 488 nm, 561 nm and 643 nm laser lines for excitation running the LASX Leica software. For image quantification, 16-bit (1024×1024) images were acquired under identical settings avoiding signal saturation. Cell and nucleus area measurements were performed by using ICY software (Quantitative Image Analysis Unit, Institut Pasteur, http://icy.bioimageanalysis.org/). A pipeline was created to completely automate image analysis by using the following sequential plugins: active contours (cell segmentation), hk-means (threshold detection), wavelet spot detector (spot detection) and studio (colocalization). Total Fluorescence Integrated intensity measurement was performed by using ImageJ (FIJI) (Schindelin et al. 2012). The number of dots (lysosomes) was performed by using the LOG detector algorithm available on the TrackMate plugin (FIJI). For live cell imaging assays, HeLa cells were grown in glass bottom culture dishes (MatTek Corporation, Ashland, MA, USA) and labeled with the following Invitrogen probes: ER-Tracker™ Blue-White DPX (E12353), LysoTracker™ Red DND-99 (L7528, Invitrogen) and Magic-red (Immunochemistry Technologies, LLC) according to the manufacturer’s protocol. Before image acquisition, culture medium was replaced with phenol red-free DMEM supplemented with HEPES (10 mM, pH 7.4), and images were acquired with the 63x oil immersion objective (1.4 NA) of the TCS SP8 laser-scanning confocal microscope, running the Leica Application Suite LAS X software, coupled to a controlled temperature chamber (UNO-temp controller, OKOLAB) acquiring 16-bit images at 37°C (488 laser for excitation; HyD: 510–550 nm; 1024 × 1024 pixels; frame average 2). For volume analysis, z-stack (0.3 om z-interval, 1024×1024, 180 om pixel size) images were quantified using 3D analysis plugin by using ICY software. Total Fluorescence Integrated intensity and dots number were measured using ImageJ.

### RNA isolation and RT-PCR analysis

Total RNA extraction from HeLa cells, oligo-dT and MMLV reverse transcriptase and quantitative reverse transcription PCR of the cDNA template (RT-PCR) was carried out as previously described (Bustamante et al. 2020). The specific primer pairs used for CYCLOPHILIN-A, CYCA (NM_001300981.2) and XBP1 (NM_001079539.2) were the following: **CYCA-F** TCGAGTTGTCCACAGTCAGC, **CYCA-R** TTCATCTGCACTGCCAAGAC, **XBP1-F** CGCTTGGGGATGGATGCCCTG and **XBP1-R** CCTGCACCTGCTGCGGACT.

### Cell viability assays

Exponentially growing cells stably expressing either FLAG-tagged HERPUD1-WT or HERPUD1-S59D were trypsinized and seeded at 20,000 cells per well in 96-well microplates and allowed to attach for 6 h at 37°C and 5% CO_2_. Then, cells were incubated with CDDP in serial dilutions from 1 mM to 1 µM in 2% FBS medium and were incubated for 24 h at 37°C and 5% CO_2_. After drug incubation, the IC50 was obtained using the Sulforhodamine B (SRB) cell cytotoxicity assay (Blois, Smith, and Josephson 2011). Briefly, cells were fixed (10% trichloroacetic acid, 4°C, 1 h), water washed, dried and stained (0.4% SRB v/v in 0.1% acetic acid, 1 h at RT), and then washed four times (1% acetic acid). Dissolved SRB (10 mM Tris-base, pH 10) was quantified (564 nm, Synergy HT BioTek reader). CDDP treatments were done in quadruplicate in at least three independent experiments. The IC50 values associated with the cytotoxic effects of CDDP were calculated using GraphPad Prism software (version 8.2; GraphPad Software, San Diego, CA, USA) using non-linear regression model and dose-response equations (log(inhibitor) vs. normalized response). After drug incubation, apoptotic cells were analyzed with the commercial kit Vybrant Apoptosis Pacific Blue-annexin V (cat#A35122, Invitrogen) using the protocol provided by the manufacturer. Briefly, cells stably expressing either FLAG-tagged HERPUD1-WT or HERPUD1-S59D were treated with 10 µM CDDP for 24 h. Cells were harvested and centrifuged at 800 x g for 5 min at room temperature and washed with cold PBS 1X. Cells were re-centrifuged, and the pellet resuspended in 100 µL annexin-binding buffer. Then, 5 µL of the annexin V conjugate was added to the cell suspension and incubated for 15 min at room temperature. After, 400 µL of annexin-binding buffer was added, gently mixed, and maintained on ice for later analysis in a BD FACSCanto II flow cytometer (Flow Cytometer Facility of Cell 4 Cell, Santiago, Chile), with a previous incubation with 3 µL of 7-Amino-Actinomycin D (7AAD) to exclude non-viable cells (included in the kit).

### Densitometric Quantification and Statistical Analysis

The amount of immunoblot signal was estimated using Image J software version 1.48v (Wayne Rasband, NIH, http://imagej.nih.gov). For each condition, protein bands were quantified from at least three independent experiments in order to ensure adequate statistical power. Data analysis was performed using Microsoft Excel 2013 for Windows (Redmond, WA, USA) or GraphPad Prism 6. Results are represented in graphs depicting the mean ± standard deviation. Statistical significance of data comparisons from two groups comparisons was determined with two-tailed unpaired Student’s T-test for parametric data. Values of P <0.05 (*), P <0.01 (**), P <0.001 (***) were regarded as statistically significant and are indicated in the figures.

## Acknowledgments

We thank the generous donation of pCI-HERPUD1-Flag and pCI-HERPUD1-ΔUBL-Flag by Doctor Koichi Kokame from National Cerebral and Cardiovascular Center, Japan. Additionally, we thank Dr. Lavandero and Dr. Quiroga for the critical discussion of the manuscript.

## Data availability statement

All the original data supporting the results is available.

## RESULTS

### HERPUD1 is a regulator of autophagy under the control of its UBL domain

Previous reports have demonstrated a close interplay between autophagy and the Ubiquitin-Proteasome System (Bustamante et al. 2018), however, to date, few proteasomal substrates are known as modulators of autophagy (Jia and Bonifacino 2019; Guarascio et al. 2020; Thayer et al. 2020). To search for potential novel candidates that could be downregulated by the proteasome in order to activate autophagy, we performed a SILAC-based proteomic study to quantitatively determine the proteome of H4 neuroglioma cells under basal and induced autophagy by EBSS starvation conditions. To eliminate all proteins downregulated because of autophagy activation, we compared the proteome of H4 cells where autophagy is inhibited by stable depletion of ATG5 by shRNA-mediated knockdown (shATG5) respect to control H4 cells expressing an shRNA against the luciferase gene (shLuc), both cell lines previously characterized (González et al. 2017; Tapia et al. 2019) (Fig. 1A). ATG5 protein is part of a complex with ATG12 and ATG16L that controls an essential step in the autophagosome formation (Walczak and Martens 2013). Silencing of ATG5 causes a strong inhibition in LC3B-positive autophagosomes, a phenotype also previously confirmed in our H4 cell lines (González et al. 2017). Among all the proteins downregulated by EBSS starvation, we found that in both H4 cell lines, shLuc (Fig. 1B) and shATG5 (Fig. 1C) the most significantly downregulated protein was HERPUD1, a protein originally identified as a homocysteine-inducible gene, that is also upregulated by endoplasmic reticulum (ER) stress (K. Kokame et al. 2000; Koichi Kokame, Kato, and Miyata 2001). Importantly, HERPUD1 is an ER-stress membrane protein whose levels under non-stressful conditions are low due to proteasome degradation (K. Kokame et al. 2000; Sai et al. 2003). Indeed, pharmacological inhibition of the proteasome leads to a rapid increase of HERPUD1 levels (Sai et al. 2003; Miura et al. 2010). In addition to HERPUD1, we found several other proteins significantly down- or up-regulated by EBSS treatment (Fig. 1D). However, while some proteins were down- or up-regulated by EBSS treatment in both, shLuc and shATG5 stable expressing cell lines (Fig. 1D, red dots), many other hits were only down or up-regulated dependent on ATG5 protein expression (Fig. 1D, purple dots). The complete list of proteins that responded significantly to EBSS treatment in both cell lines is shown in Suppl. Fig 1. Here, we focus on the characterization of HERPUD1, the hit with the highest score of downregulation in both cell lines, working with the hypothesis that its reduction by EBSS treatment could be indicative of its role as negative regulator of autophagy activity. Previous studies had shown that silencing HERPUD1 triggers the autophagic flux (Quiroga et al. 2013). Therefore, in this study, and considering that HERPUD1 levels are increased under ER stress, we study the cellular consequences of HERPUD1 stability on autophagy function. HERPUD1 is an integral membrane protein with both termini facing the cytoplasm, with the ubiquitin-like domain (UBL) located in its N-terminus (Fig. 2A; UBL: purple color, left side). Here, we cloned full-length HERPUD1 into the pLVX-IRES-Puro vector with a FLAG-tagged in its C-terminal end (Fig. 2A; FLAG: green color) designed as HERPUD1-WT. In addition, we cloned a FLAG-tagged HERPUD1 version lacking the residues between Val^10^-Cys^86^ designed as HERPUD1-ΔUBL (Fig. 2A; right side), as previously reported (Sai et al. 2003). Further, we generated puromycin-resistant HeLa cells stably expressing either HERPUD1-WT or HERPUD1-ΔUBL, considering that previous characterization of HERPUD1 in HeLa cells was done only by transient transfection (Sai et al. 2003). Western blot analysis using anti-HERPUD1, and anti-FLAG antibodies showed that the levels of HERPUD1-ΔUBL were higher than the levels induced by the overexpression of HERPUD1-WT (Fig. 2B, lane 3 and lane 5). In addition, western blot analysis with an anti-HERPUD1 antibody confirmed HERPUD1 is expressed in low levels under non-stressful conditions (Fig. 2B, lane 1). The UBL is a domain that resembles ubiquitin in terms of their primary sequence and three-dimensional structure, considered a general interaction motif with the proteasome particularly with the 19S regulatory particle of the 26S proteasome (R. Hartmann-Petersen and Gordon 2004). Interestingly, it has been previously shown that deletion of UBL in HERPUD1 abolishes its proteasomal degradation (Sai et al. 2003). To confirm this, we treated the cells during 4 h with 20 µM MG132, a potent blocker of the proteasome activity (Tsubuki et al. 1996; D. H. Lee 1998). As expected, we observed that this treatment caused an increase in the levels of overexpressed HERPUD1-WT (Fig. 2B, lane 3 and 4). Similarly, we observed a slight increase in the levels of endogenous HERPUD1 (Fig. 2B, lane 1 and 2). In contrast, HERPUD1-ΔUBL did not respond to this treatment, observing similar levels in the absence or presence of MG132 (Fig. 2B, lane 5 and 6). Together, these results confirmed HERPUD1-ΔUBL is a tool to explore the effect of HERPUD1 stability on autophagy.

**Figure 2.**
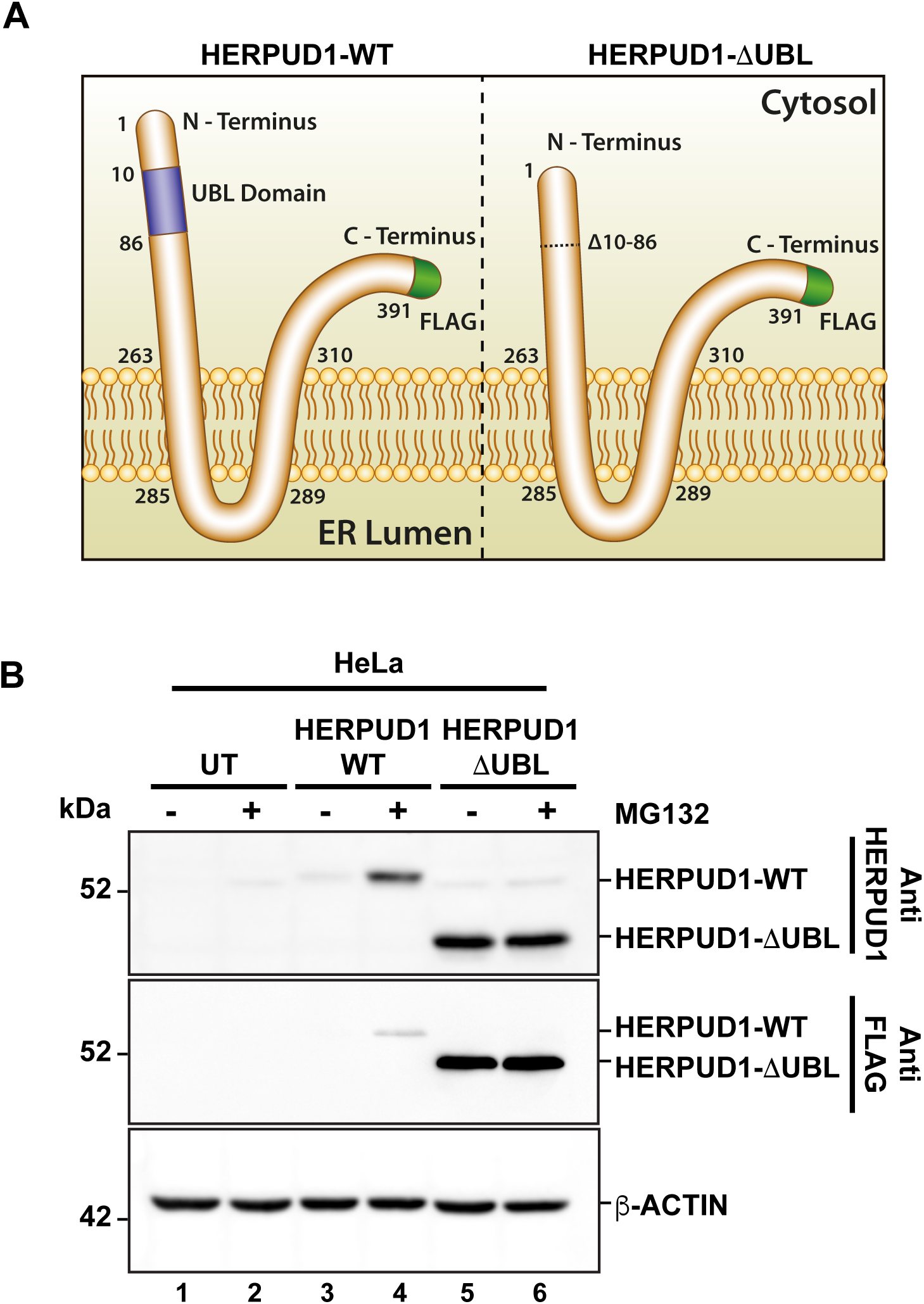
Deletion of UBL domain increases HERPUD1 protein levels. **(A)** Schematic representation HERPUD1-WT (left image) and ΔUBL (right image) at the ER membrane, both FLAG-tagged at the carboxyl-terminus (shown in green). The Ubiquitin-Like domain (UBL, amino acids 10-86) are represented in purple. **(B)** HeLa cells untransfected (UT) or stably expressing the WT or ΔUBL versions of HERPUD1 were not treated (lanes 1, 3 and 5) or treated with 20 µM of MG132 for 4 h (lanes 2, 4 and 6). Detergent-soluble protein extracts were analyzed by western blot with anti-HERPUD1 and anti-FLAG antibodies. β-ACTIN was used as loading control. Position of molecular mass markers is indicated on the left.

Therefore, we investigated the effect of HERPUD1-ΔUBL stable expression by analyzing the subcellular distribution of the microtubule-associated protein 1 light chain 3B (LC3B), a classical marker of autophagy, from here referred for simplicity as LC3 (Tanida, Ueno, and Kominami 2008). Immunofluorescence analysis of LC3 in cells expressing HERPUD1-WT showed basal autophagy represented by LC3-positive membrane dots that correspond to autophagosomes decorated with lipidated LC3 (LC3-II) (Fig. 3A; left, control). In comparison, we noticed that HERPUD1-ΔUBL cells showed a slight decrease in the number of autophagosomes which was accompanied with a higher LC3 cytosolic distribution (LC3-I) (Fig. 3A; right, control). Because autophagosomes are constantly forming autolysosomes through the fusion with acidic lysosomes for degradation, we tested the effect of BafA1, a drug that raises the lysosomal pH resulting in the perturbation of the autophagic flux (González et al. 2017). As expected, treatment of HERPUD1-WT cells with BafA1 showed a higher number of autophagosomes (Fig. 3A, left, BafA1). In contrast, this treatment in HERPUD1-ΔUBL cells showed fewer numbers of autophagosomes compared to HERPUD1-WT cells (Fig. 3A, right, BafA1). Furthermore, we biochemically validated these results performing western blot analysis of endogenous LC3 (Fig. 3B). We found that under basal conditions overexpression of HERPUD1-ΔUBL caused an increase in LC3-I levels, in comparison to HERPUD1-WT (Fig. 3B, lane 1 and 2). In agreement with this observation, HERPUD1-ΔUBL cells showed a significant reduction in the LC3-II/LC3-I ratio (0.47 ± 0.22), respect to control cells (1.00 ± 0.05) (Fig. 3C). A similar finding was observed in the presence of BafA1 treatment, observing in HERPUD1-ΔUBL cells a significant decrease in the ratio LC3-II/LC3-I (0.94 ± 0.37), respect control cells (3.01 ± 0.78) (Fig. 3B, lane 3 and 4). Importantly, similar findings were obtained under induced autophagy by EBSS starvation medium (Suppl. Fig. 2). We observed that the LC3-II/LC3-I ratio in HERPUD1-ΔUBL cells in the presence of EBSS plus BafA1 was diminished with respect HERPUD1-WT cells (Suppl. Fig. 2A, line 5 and 6 and Suppl. Fig. 2B). Altogether, these findings strongly indicate that increased stability of HERPUD1 plays a negative effect on autophagy.

**Figure 3.**
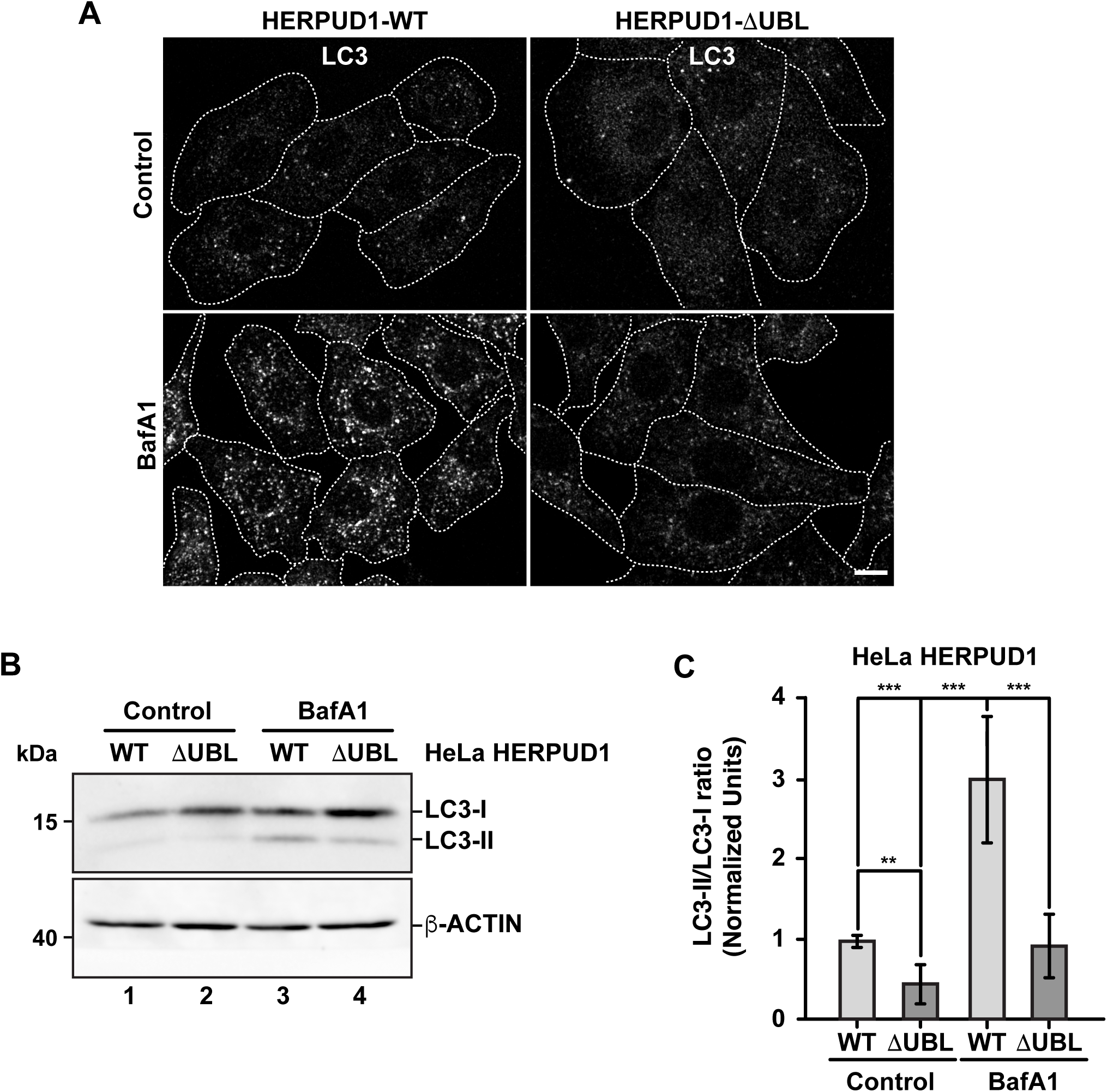
The stabilization of HERPUD1 by its UBL deletion negatively regulates autophagy. **(A)** HeLa cells stably expressing HERPUD1-WT-FLAG or HERPUD1-ΔUBL-FLAG were grown in glass coverslips and were treated or not with 100 nM BafA1 for 4 h. Cells were fixed and incubated with an antibody to LC3 followed by incubation with Alexa-488-conjugated donkey anti-rabbit IgG. Stained cells were examined by confocal microscopy. Scale bar 10 µm. **(B)** HeLa cells stably expressing HERPUD1-WT-FLAG or HERPUD1-ΔUBL-FLAG untreated (lanes 1 and 2) or treated with 100 nM BafA1 for 4 h (lanes 3 and 4). Detergent-soluble protein extracts were analyzed by western blot with a rabbit polyclonal antibody against LC3. Monoclonal antibody against β-ACTIN (clone BA3R) was used as loading control. Position of molecular mass markers is indicated on the left. **(C)** Densitometry quantification of LC3-I and LC3-II protein levels from images as those shown in B. Relative levels are expressed as the ratio of LC3-II to LC3-I. Bars represent the mean ± standard deviation. Statistical analysis was performed using two-tailed unpaired Student’s t-test (n = 3 ** p < 0.01, *** p < 0.001)

### Increased stability of HERPUD1 by the deletion of its UBL domain triggers expansion of the ER organized in stacked tubules and crystalloid ER-like structures in the absence of ER stress

Because previous reports have shown that defective autophagy leads to ER expansion (Ou et al. 1995; Jung et al. 2008) we investigated whether HERPUD1 stability could be linked with ER remodeling. In contrast to previous studies where only transient expression of HERPUD1-ΔUBL was characterized (Sai et al. 2003), we investigated the subcellular distribution of FLAG-tagged HERPUD1-ΔUBL but in stable HeLa cell lines, in comparison with its wild type version. Immunofluorescence analysis with an anti-FLAG antibody showed a stronger signal for HERPUD1 in the absence of its UBL domain (Fig. 4A, FLAG, lower left panel) in comparison to HERPUD1-WT (Fig. 4A, upper left panel). A similar result was also observed with an anti-HERPUD1 antibody (Suppl. Fig. 3A). To unveil if this phenotype was affecting the ER in general, we analyzed by immunofluorescence the distribution of endogenous ER proteins markers. First, we tested endogenous CALNEXIN, an ER membrane resident protein that acts as a molecular chaperone of glycoproteins (Ou et al. 1995) and GRP94, an ER resident membrane protein of the Heat-Shock Protein (HSP) 90 family (Marzec, Eletto, and Argon 2012). We found profound changes in the distribution of both proteins showing a similar pattern to anti-FLAG in HERPUD1-ΔUBL cells (Fig. 4A, CALNEXIN and GRP94, lower panel), compared to HERPUD1-WT where a characteristic ER pattern is observed. Indeed, HERPUD1-ΔUBL shows a nice colocalization with CALNEXIN and GRP94 (Fig. 4A, lower merge). In addition, and to exclude the possibility that this phenotype was specific to CALNEXIN and GRP94, we analyzed the ER pattern using an ER-tracker probe in live cells, observing clear changes in the morphology of the ER respect to HERPUD1-WT (Suppl. Fig. 3B). We noticed that in comparison to HERPUD1-ΔUBL, HERPUD1-WT did not show a high colocalization with CALNEXIN and GRP94 (Fig. 4A, upper merge), which can be explained by the low levels of HERPUD1-WT expression due to its constant degradation by the proteasome under basal conditions (Sai et al. 2003). In this regard, localization of HERPUD1 at the ER has been well documented (K. Kokame et al. 2000). Whereas HERPUD1-ΔUBL did not cause an increase in the levels of CALNEXIN and GRP94 measured by Western blot analysis (Suppl. Fig. 3C y 3D), we concluded that HERPUD1 stability by the deletion of its UBL domain causes an ER-like expansion phenomena that does not involve the increase in the levels of CALNEXIN and GRP94. HERPUD1 is known to be upregulated under ER stress, a condition reported to cause ER membrane expansion to alleviate this condition (Schuck et al. 2009). Therefore, we studied whether HERPUD1-ΔUBL could be causing ER proliferation. To assess this, we performed immunofluorescence analysis with an anti-GRP94 antibody. Interestingly, we observed that HERPUD1-ΔUBL expressing cells present a large and dense ER network extending throughout the entire cytoplasm including the periphery of the cell, compared to the less extended ER network observed in HERPUD1-WT cells (Fig. 4B, left and right panel). To gain a more comprehensive understanding of the ER differences between HERPUD1-WT and HERPUD1-ΔUBL expressing cells, we measured the ER volume using 3D images at high magnification taken with a z-interval of 0.3 µm followed by a z-stack maximum intensity projection. For visualization, we analyzed the distribution of GRP94 in the whole cell using a heat map gradient ranging from non-intensity (black color) to the higher intensity (red color) (Fig. 4B, left and right panel). This analysis confirmed HERPUD1-ΔUBL increases the volume of the ER and the expansion of this organelle, compared to the expression of HERPUD1-WT. Indeed, quantitative analysis showed a significant increase in the ER volume with the expression of HERPUD1-ΔUBL (2361 ± 967), compared to HERPUD1-WT (1700 ± 689) (Fig. 4C). The ER morphological changes observed by confocal microscopy prompted us to perform ultra-structural analysis by transmission electron microscopy (TEM). While cells expressing HERPUD1-WT showed the common ER cytoplasmic structures (Fig. 4D left panel [ER]), unexpectedly HERPUD1-ΔUBL showed a high number of staked tubular ER structures oriented in a hexagonal spatial distribution (Fig. 4D center panel [ER]). Others have referred to these structures as energetically stable structures formed by a remarkable proliferation of smooth ER, resembling crystalloid structures of ER (Chin et al. 1982; Anderson, Orci, and Brown 1983; Borgese, Francolini, and Snapp 2006), which in appearance can be compared with “honeycomb like-structures” (Fig. 4D center panel [ER]). Additionally, a higher magnification (Fig. 4D, right panel), confirmed the ER contained in the crystalloid structure corresponds to smooth ER evidenced by the absence of attached ribosome structures.

**Figure 4.**
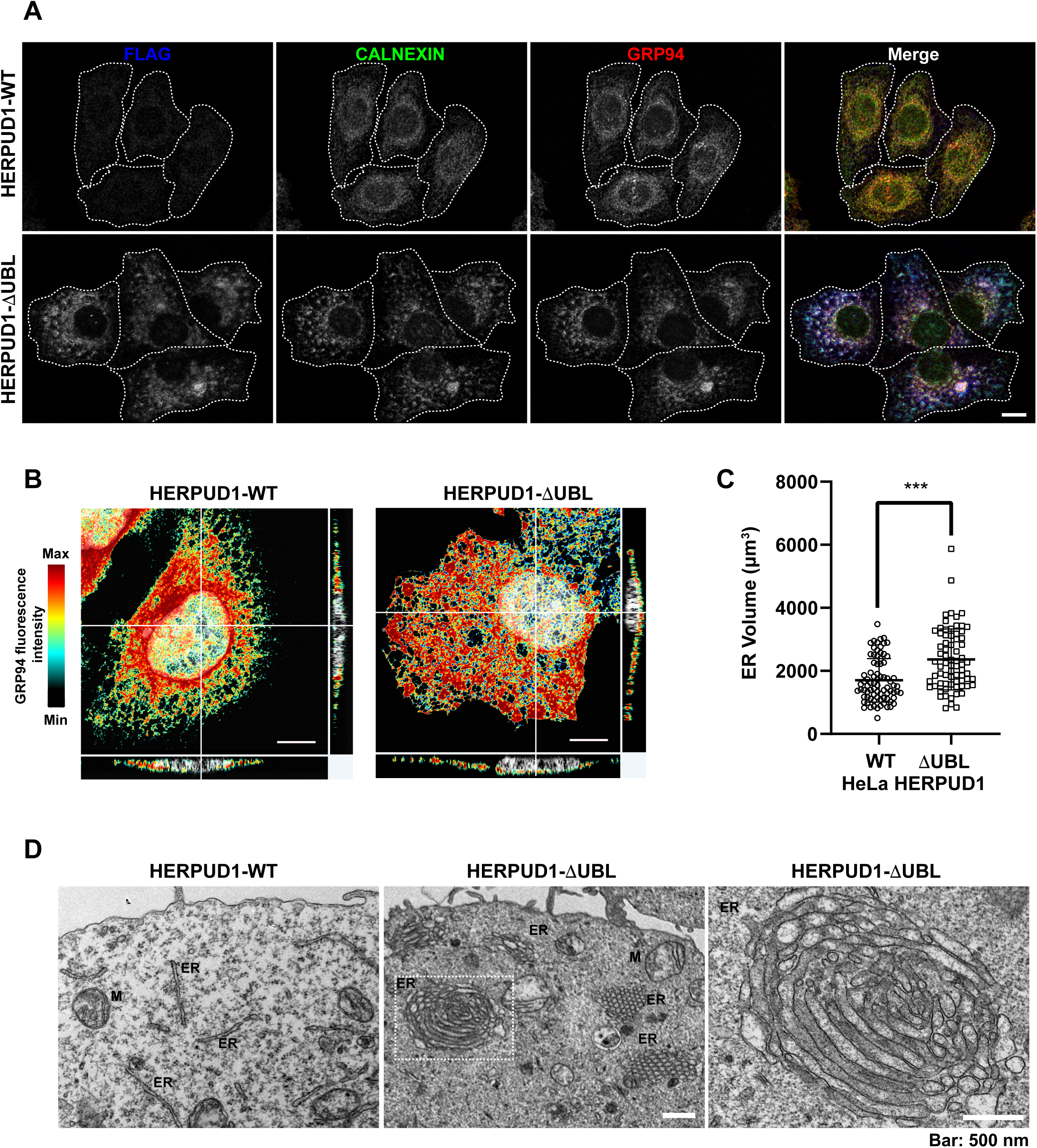
HERPUD1-ΔUBL stabilization alters the ER morphology. **(A)** HeLa cells stably expressing HERPUD1-WT-FLAG or HERPUD1-ΔUBL-FLAG grown in glass coverslips were fixed, permeabilized and triple-labeled with a mouse monoclonal antibody against FLAG, with a rabbit monoclonal antibody against CALNEXIN and with a rat monoclonal antibody against GRP94 followed by incubation with Alexa-488-conjugated donkey anti-rabbit IgG (green channel), Alexa-594-conjugated donkey anti-rat IgG (red channel) and Alexa-647-conjugated donkey anti-mouse IgG (blue channel). Images were acquired using a TCS SP8 laser-scanning confocal microscope. The fourth image on each row is the merge of blue, green and red channels; yellow indicates colocalization of the red and green channels, cyan indicates colocalization of the green and blue channels, magenta indicates colocalization of the red and blue channels, and white indicates colocalization of all three channels. Scale bar, 10 µm. **(B)** HeLa cells stably expressing HERPUD1-WT-FLAG or HERPUD1-ΔUBL-FLAG grown in glass coverslips were fixed, permeabilized and labeled with rat monoclonal antibody against GRP94 followed by incubation with Alexa-594-conjugated donkey anti-rat IgG. Stained cells were examined by fluorescence microscopy. Images were acquired using a TCS SP8 laser-scanning confocal microscope. Pseudocolor image was created using all the serial confocal sections of HeLa HERPUD1-WT-FLAG (left image) and HERPUD1-ΔUBL-FLAG (right image). The scale on the right represents the maximum (red) to minimum (black) intensity measured for GPR94. **(C)** Quantification of ER-volume obtained from serial image reconstruction (z-stack 0.3 µm z-interval, 1024×1024, 180 µm pixel size). Volume is depicted in a scatter plot, open circles represent HeLa HERPUD1-WT-FLAG (n=79) and open squares HERPUD1-ΔUBL-FLAG (n=72). Statistical analysis was performed using two-tailed unpaired Student’s t-test (*** p < 0.001). **(D)** TEM micrograph shows HeLa HERPUD1-WT-FLAG or HERPUD1-ΔUBL-FLAG (left and center images, respectively) at a lower magnification. Crystalloid ER structure is visible as a honeycomb in HERPUD1-ΔUBL (central image). An ER crystalloid structure from HERPUD1-ΔUBL-FLAG at a higher magnification is shown (right image from dashed square in the center image, ER: endoplasmic reticulum; M: mitochondria). Scale bar 500 nm.

To gain insights whether the ER proliferation triggered by the expression of HERPUD1-ΔUBL was the result of an ER stress response, we studied X-box binding protein 1 (XBP1) as a reporter of ER stress and the unfolded protein response (UPR). First, we analyzed by RT-PCR the XBP1 mRNA processing, from the unspliced inactive XBP1 mRNA (u*XBP1*) form to the spliced active XBP1 mRNA (s*XBP1*) form (Fig. 5A), which is considered a hallmark of the UPR response (Yoshida et al. 2001). RT-PCR analysis of mRNA from HeLa WT untreated cells showed a single band as expected (Fig. 5A, lane 1). The same analysis including tunicamycin (Tun) (inhibitor of N-linked glycosylation (Heifetz, Keenan, and Elbein 1979) and thapsigargin (Tg) (blocker of ER Ca2+ import (Thastrup et al. 1990)) as positive inducers of ER stress, produced the appearance of two bands in HeLa-WT cells, indicating the s*XBP1* form in response to ER stress (Fig. 5A, lane 2 and 3). In contrast, a stable expression of either HERPUD1-WT or HERPUD1-ΔUBL showed no detection of s*XBP1,* observing only the expression of the u*XBP1* form (Fig. 5A, lane 4 and 5), similar to control HeLa WT untreated cells (Fig. 5A, lane 1). Analysis of PCR bands relative to *CYCLOPHILIN A (CYC-A)* as a housekeeping control showed no difference between cells expressing either HERPUD1-WT or HERPUD1-ΔUBL proteins (Fig. 5A, lane 4 and 5). Similar findings were observed by western blot analysis (Fig. 5B). s*XBP1* is rapidly translated to a highly active transcription factor, known as XBP1, responsible for the upregulation of a variety of UPR genes (Yoshida et al. 2001). To examine the significance of the UBL domain of HERPUD1 in the response to ER stress induction, HERPUD1-WT cells and HERPUD1-ΔUBL cells were treated with 2 μM Tg for 2, 4 and 6 h and we performed western blot analysis for XBP1 detection. We observed XBP1 protein was almost undetectable in both cell lines in the absence of a stressor (Fig. 5B, lane 1 and 5). In contrast, we observed a robust induction of XBP1 in both cell lines upon treatment for 2, 4 and 6 h with 2 μM Tg, observing similar XBP1 expression levels after 6 h of treatment in both cell lines (Fig. 5B, lane 4 and 8). Quantitative analysis confirmed this conclusion, observing no significant differences between HERPUD1 WT (58.99 ± 9.18) and HERPUD1-ΔUBL (59.59 ± 17.49) expressing cells (Fig. 5C). In addition to XBP1, we tested by western blot analysis the ER resident transmembrane protein PERK, a kinase that undergoes hyperphosphorylation in response to ER stress shown as a shift in its mobility during SDS-polyacrylamide gel electrophoresis (Bertolotti et al. 2000). This kinase mediates the attenuation of the global translation by the phosphorylation of eIF2ɑ (Bertolotti et al. 2000; Harding, Zhang, et al. 2000). In contrast to its differential electrophoretic migration with Tun and Tg (Suppl. Fig. 4, lane 2 and 3), no changes in PERK migration were observed in HERPUD1-WT or HERPUD1-ΔUBL cells (Suppl Fig. 4, lane 4 and 5). Analysis of the activating transcription factor 4 (ATF4), a protein highly expressed upon UPR response (Harding, Novoa, et al. 2000) showed again no changes in ATF4 levels in either cell line (Suppl. Fig. 4, lane 4 and 5). In contrast, a robust detection of ATF4 was found upon the addition of Tun and Tg (Suppl. Fig. 4, lane 2 and 3). Altogether, our findings strongly indicate that the maintenance of the ER expansion triggered by HERPUD1 stability is not the consequence of ER stress.

**Figure 5.**
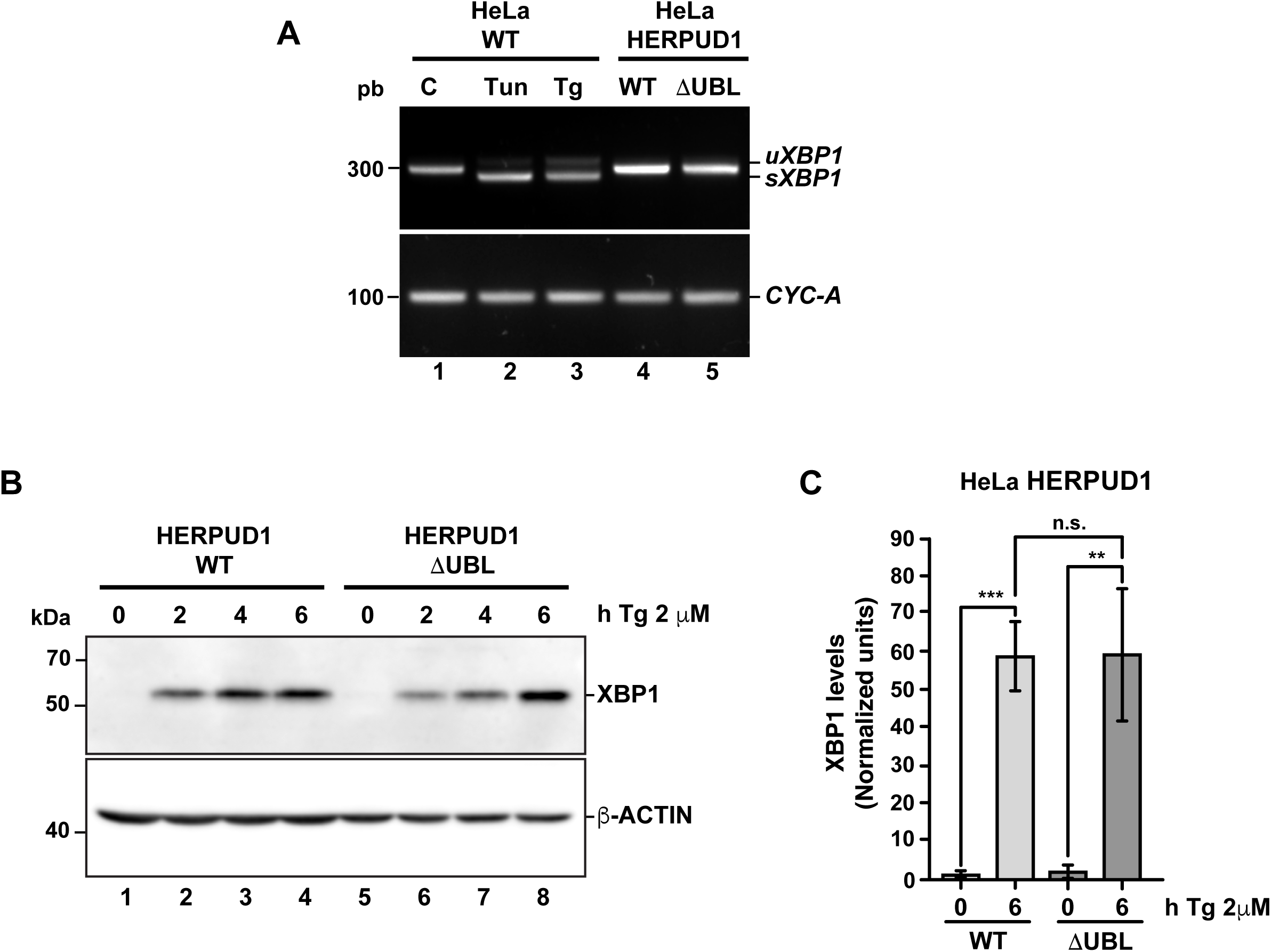
The stability of HERPUD1 by the deletion of its UBL domain does not trigger endoplasmic reticulum stress. **(A)** Splicing of *XBP1* was analyzed by RT-PCR. Total RNAs were obtained from HeLa cells stably expressing HERPUD1-WT-FLAG or HERPUD1-ΔUBL-FLAG, additionally, total RNA was obtained from HeLa WT cells untreated or treated with 2 µM Thapsigargin (Tg) or 5 µg/mL Tunicamycin (Tun) for 4 h as a control of XBP1 splicing. Then, cDNA was synthesized and mRNA expression of XBP1 was analyzed using specific primers. (*uXBP1*: unspliced form, *sXBP1*: spliced form). CYCLOPHILIN-A (CYC-A) was used as an internal control. **(B)** HeLa cells stably expressing HERPUD1-WT-FLAG or HERPUD1-ΔUBL-FLAG were treated with 2 µM Thapsigargin (Tg) for different timepoints (0 h, 2 h, 4 h and 6 h). Detergent-soluble protein extracts were analyzed by western blot with a monoclonal antibody against XBP1. Monoclonal antibody against β-ACTIN was used as loading control. Position of molecular mass markers is indicated on the left. **(C)** Densitometry quantification of XBP1 protein levels from images as those shown in B. Bars represent the mean ± standard deviation. Statistical analysis was performed using two-tailed unpaired Student’s t-test (n = 3 n.s not statistically significant, ** p < 0.01, *** p < 0.001).

### Increased HERPUD1 stability by the deletion of its UBL domain leads to a remarkable increase in lysosomal number and function

ER proliferation that excludes ribosomes is able to regulate ER contacts with lysosomes (Friedman et al. 2013; C. A. Lee and Blackstone 2020). Lysosomes are an important catabolic organelle of eukaryotic cells, containing a diverse repertoire of acidic hydrolases (luminal pH of 4.5–5.0) that can digest macromolecules such as sugars, lipids and proteins and even entire organelles (H. Xu and Ren 2015). We asked whether expanded ER by increased HERPUD1 stability could have an impact on endolysosomal organelles. To visualize these compartments, HERPUD1-WT and HERPUD1-ΔUBL expressing HeLa cells were stained with an antibody against the endogenous lysosomal-associated membrane protein 1 (LAMP1), a membrane protein found on late endosomes and lysosomes (Griffiths et al. 1988). Unexpectedly, we observed a robust increase in LAMP1 positive structures located around the entire cytoplasm (Fig. 6A). In agreement with this finding, quantification analysis showed a significant increase in the number of LAMP1 positive structures in HERPUD1-ΔUBL (88.72 ± 16.77) respect to control cells (69.03 ± 13.79) (Fig.6B). To determine if the increase in LAMP1 structures corresponded to lysosome organelles, we performed co-staining of LAMP1 with CATHEPSIN-D (CAT-D), a luminal hydrolytic lysosomal enzyme (Benes, Vetvicka, and Fusek 2008), observing several LAMP1 positive structures positive to CAT-D (Suppl. Fig 5A, lower merge). In addition, we noticed a significant increase in the integrated fluorescence intensity of CAT-D in HERPUD1-ΔUBL (1.47 ± 0.88), compared to HERPUD1-WT expressing cells (1.00 ± 0.20) (Suppl. Fig 5B). Considering this increase in CAT-D fluorescence we evaluated if HERPUD1-ΔUBL also impacts lysosomal activity, as assessed by measuring lysosomal acidity and activity in live cells. We first measured if HERPUD1 stability affects the range of lysosomal pH, using the pH-sensitive lysosomal dye LysoTracker Red, which accumulates and emits red fluorescence in acidic compartments with pH <6.5 (De Duve et al. 1974; Chou, Paul Krapcho, and Hacker 2001). Then, we measured CATHEPSIN-B activity using the Magic Red assay (Boonacker et al. 2003). As shown in Figure 6C, expression of HERPUD1-ΔUBL causes a significant increase in the number of LysoTracker positive structures (211.95 ± 78.90), compared to control cells (117.85 ± 48.97) (Fig. 6C and 6D). Similarly, we found an increase in the number of structures positive to Magic Red fluorescence (Figure 6E). This was confirmed by measuring the integrated fluorescence intensity, which revealed a significant increase of this parameter in HERPUD1-ΔUBL cells (1.56 ± 0.43), compared to control cells (1.00 ± 0.19) (Fig. 6F). Altogether, these findings show that ER expansion triggered by HERPUD1 increased stability is a potent strategy to promote lysosomal function.

**Figure 6.**
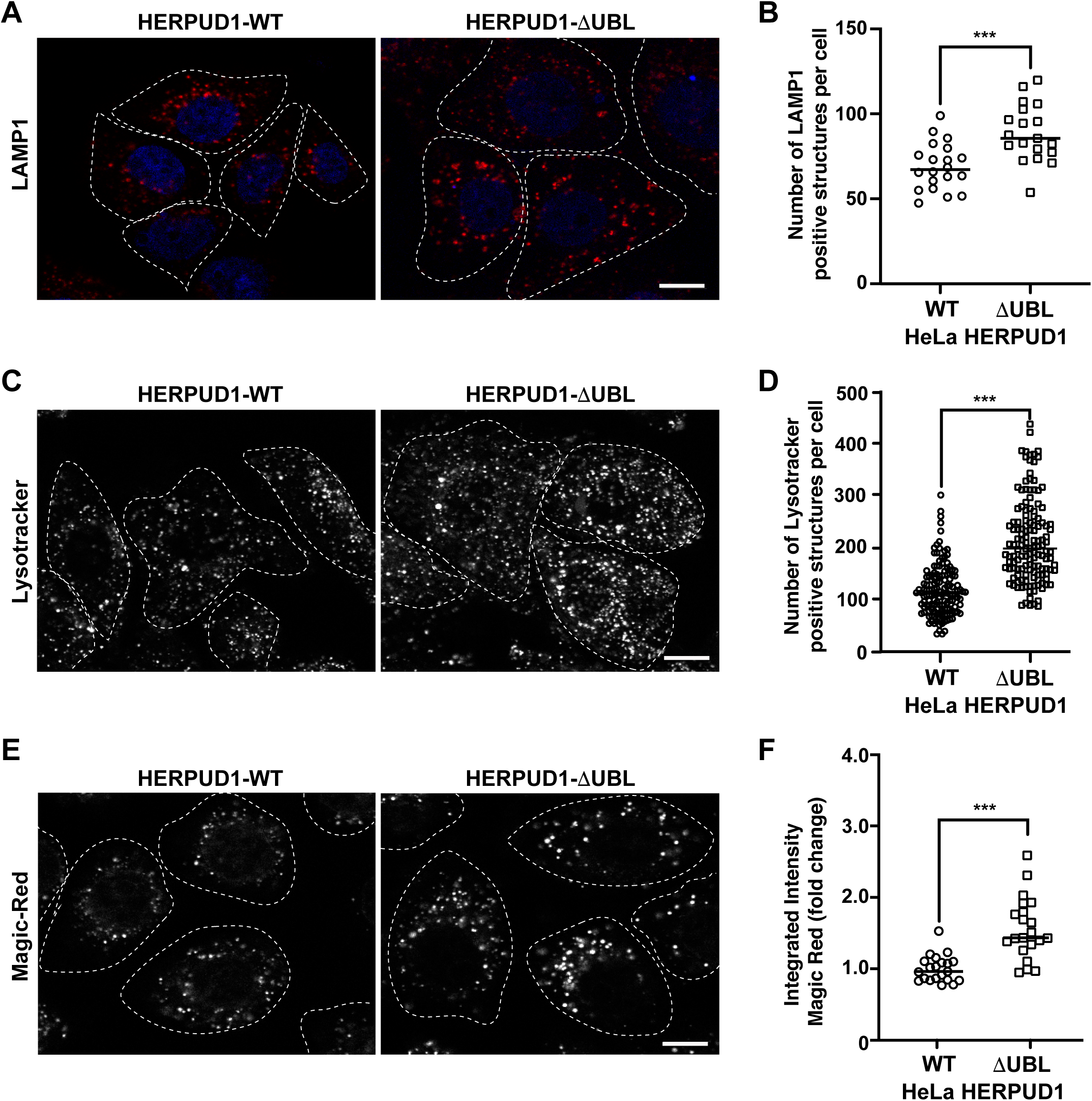
Expression of HERPUD1-ΔUBL increases lysosomal number and function. **(A)** HeLa cells stably expressing HERPUD1-WT-FLAG or HERPUD1-ΔUBL-FLAG grown in glass coverslips were fixed, permeabilized and labeled with a mouse monoclonal antibody against LAMP1 followed by incubation with Alexa-594-conjugated donkey anti-mouse IgG, and DAPI for nuclei staining. Stained cells were examined by fluorescence microscopy. Images were acquired using a TCS SP8 laser-scanning confocal microscope. Scale bar 10 µm. **(B)** Quantification of LAMP1 positive structures per cell. In the scatter plot each circle and square represents the average of positive LAMP1 dots per cell from a frame of HeLa HERPUD1-WT (n=559) or HERPUD1-ΔUBL (n=700) respectively. The quantified cells were from 3 independent experiments. **(C and E)** HeLa cells stably expressing HERPUD1-WT-FLAG or HERPUD1-ΔUBL-FLAG were grown in glass bottom culture dishes and labeled with **(C)** LysoTracker™ Red DND-99 or **(E)** Magic Red®. For live cell imaging analysis, culture medium was replaced with phenol red-free DMEM supplemented with HEPES (10 mM, pH 7.4) and images were acquired with TCS SP8 laser-scanning confocal microscope at 37°C. Scale bar 10 µm**. (D)** Quantification of LysoTracker™ dots per cell. Open circles and open squares represent the average of fluorescence signals for each cell of HeLa HERPUD1-WT-FLAG (n=139) or HERPUD1-ΔUBL-FLAG (n=138) respectively. **(F)** Quantification of Magic Red® positive structures. In the scatter plot each circle and square represents the average of positive dots for Magic Red per cell from a frame of HeLa HERPUD1-WT-FLAG (n=1279) or HERPUD1-ΔUBL-FLAG (n=984) respectively. The quantified cells were from 3 independent experiments. Statistical analysis was performed using two-tailed unpaired Student’s t-test (*** p < 0.001)

### The phosphomimetic mutant S59D in the UBL domain promotes HERPUD1 stability mimicking the effects of UBL deletion on the ER and autophagy

Further, we searched for a mechanism that could mimic the phenotype observed by the deletion of UBL on HERPUD1. As shown by the analysis with PyMOL molecular graphics system (Suppl. Fig 6A), the UBL domain of HERPUD1 (PDB 1WGD; green color) resembles UBIQUITIN (PDB 2MSG; purple color) in terms of their three-dimensional structure. UBIQUITIN as well as UBL domains are known targets of phosphorylation under cellular stress (Kondapalli et al. 2012; Swaney, Rodríguez-Mias, and Villén 2015; Sauvé et al. 2018). In fact, oxidative stress promotes phosphorylation of UBIQUITIN at Ser65 causing the accumulation of ubiquitylated proteins due to a reduction in global protein turnover rates (Koyano et al. 2014; Swaney, Rodríguez-Mias, and Villén 2015). In the same line, it is known that the UBL domain on PARKIN is phosphorylated by PINK1, and is responsible for the activation of its E3 ubiquitin-ligase activity (Sauvé et al. 2018), both essential players of mitochondria quality control by mitophagy (Jin and Youle 2012). Thereby, we searched for Ser residues as possible candidates of phosphorylation in the UBL domain of HERPUD1 using the KinasePhos2.0 web tool. Five residues were predicted in position Ser16, Ser27, Ser33, Ser59 and Ser90 (Suppl. Fig 6B, Ser residues in red color). For all these five Ser residues, we generated phosphoinert (substitutions to alanine) and phosphomimetic (substitutions to aspartic acid) mutant versions. Stably transfected HeLa cells were generated for each mutant. Among all mutations tested, the substitution S59D, but not the S59A, showed the strongest increase in HERPUD1 signal, as is shown by immunofluorescence with an anti-FLAG antibody (Fig. 7A and 7B). Importantly, we found that the phosphomimetic S59D mutant presents an ER pattern very similar to HERPUD1-ΔUBL cells, confirmed by the co-staining with GRP94 (Fig. 7A, right panel merge). In contrast, no changes were observed with the phosphoinert S59A mutant, showing a similar phenotype than HERPUD1-WT (Fig. 7A and 7B). Thus, further characterization included the comparison between HERPUD1-WT and HERPUD1-S59D. In addition, under basal conditions western blot analysis using anti-HERPUD1 and anti-FLAG antibodies showed higher levels of HERPUD1-S59D (40.09 ± 5.31) compared to either HERPUD1-WT (1.00 ± 0.53) or HERPUD1-S59A (2.88 ±1.76) (Fig. 7C, lane 5 compared to lane 1 and 3 and Fig. 7D). We also measured the levels of these versions upon treatment for 4 h with 20 mM MG132, which is a proteasomal inhibitor, known to abolish HERPUD1 proteasomal degradation (Yan et al. 2014). As expected, MG132 treatment caused a significant increase in HERPUD1-WT (16.55 ± 7.13) (Fig. 7C, lane 1 compared to lane 2 and Fig. 7D). A similar result was obtained with HERPUD1-S59A (20.36 ± 9.65) (Fig. 7C, lane 3 compared to lane 4 and Fig. 7D). In contrast, we observed non-significant differences in the levels of HERPUD1-S59D in the absence (40.09 ± 5.31) or presence of MG132 (47.17 ± 11.74) (Fig. 7C, lane 5 and 6 and Fig. 7D). Further, and based on the negative impact of HERPUD1 stability on autophagy by the deletion of its UBL domain, we next investigated if the stability of HERPUD1-S59D could have a similar outcome. Biochemically, we found that HERPUD1-S59D cells have a significant reduction in the LC3-II/LC3-I ratio (0.66 ± 0.02), in comparison to HERPUD1-WT (1.00 ± 0.18) (Fig. 7E and 7F). This result was corroborated by immunofluorescence analysis of LC3 in the absence or presence of BafA1 showing that HERPUD1-S59D cells present a lesser number of autophagosomes, compared to HERPUD1-WT (Suppl. Fig. 7). Moreover, in agreement with previous reports indicating that a reduction in autophagy enhances ER expansion and cell size (Khaminets et al. 2015; Miettinen and Björklund 2015), we noticed that HERPUD1-S59D cells were significantly larger in size (2.52 ± 0.98), compared to HERPUD1-WT (1.00 ± 0.4) and HERPUD1-S59A cells (0.88 ± 0.29) (Fig. 7A and Suppl. Fig. 8A). Importantly, cell size quantification showed HERPUD1-ΔUBL were also larger in size (1.36 ± 0.64) than HERPUD1-WT (1.00 ± 0.4), but smaller than HERPUD1-S59D (2.52 ± 0.98) (Suppl. Fig. 8A). In addition, because nuclei size remains proportional to the cell size in a wide range of genetic backgrounds and growth conditions (Huber and Gerace 2007), we studied the nuclei size in all cell lines. Surprisingly, nuclei size in HERPUD1-S59D cells were significantly larger (1.74 ± 0.72) in comparison to all tested cell lines; HERPUD1-WT (1.00 ± 0.49), HERPUD1-S59A (1.12 ± 0.47) and HERPUD1-ΔUBL (1.11 ± 0.52) (Suppl. Fig. 8B). Altogether, our findings support the idea that HERPUD1 increased stability might play an important function in cellular plasticity by commanding a program that controls ER expansion with impact in cell and nuclei size. The results with the phosphomimetic version of HERPUD1 opens the alternative of a program activated under the control of phosphorylation.

**Figure 7.**
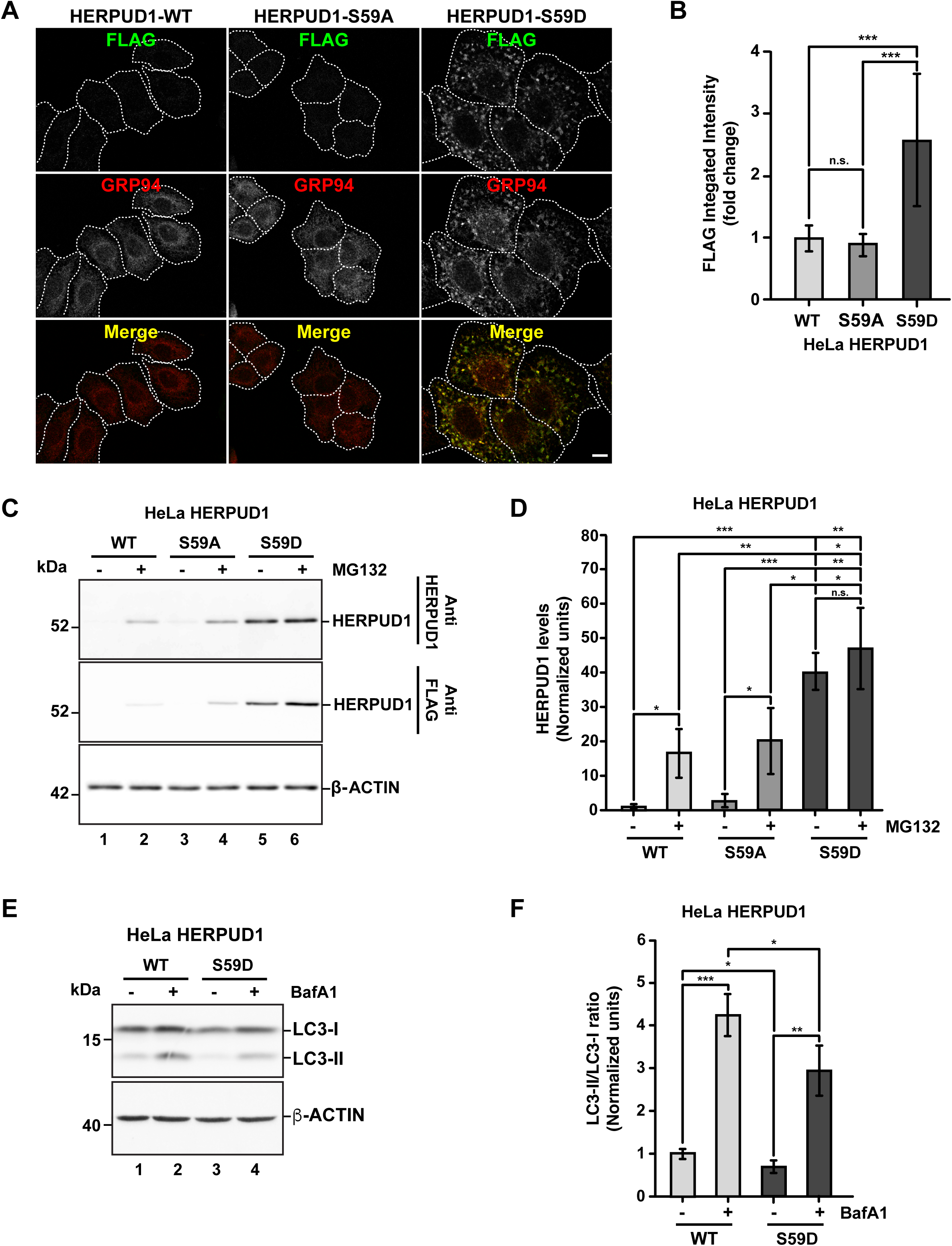
Stabilization of HERPUD1 by S59D mutation alters the ER morphology and decreases autophagy. **(A)** HeLa cells stably expressing HERPUD1-WT-FLAG, HERPUD1-S59A-FLAG or HERPUD1-S59D-FLAG were grown in glass coverslips, fixed, permeabilized, and double-labeled with mouse monoclonal antibody against FLAG and with a rat monoclonal antibody against GRP94 followed by incubation with Alexa-488-conjugated donkey anti-rabbit IgG (green channel) and Alexa-594-conjugated donkey anti-rat IgG (red channel). Images were acquired using a TCS SP8 laser-scanning confocal microscope. The third image on each column is the merge of green and red channels; yellow indicates colocalization of red and green channels. Scale bar: 10 µm. **(B)** Quantification of fluorescence FLAG signal from indicated cells. **(C)** HeLa cells stably expressing HERPUD1-WT-FLAG, HERPUD1-S59A-FLAG or HERPUD1-S59D-FLAG were treated or not with 10 µM of MG132 for 4 h. Detergent-soluble protein extracts were analyzed by western blot with anti-HERPUD1 and anti-FLAG antibodies. β-ACTIN was used as a loading control. Position of molecular mass markers is indicated on the left. **(D)** Densitometry quantification of HERPUD1 western blot signal from images as those shown in C. Bars represent the mean ± standard deviation. **(E)** HeLa cells stably expressing HERPUD1-WT-FLAG or HERPUD1-S59D-FLAG untreated (lanes 1 and 3) or treated with 100 nM BafA1 for 4 h (lanes 2 and 4). Detergent-soluble protein extracts were analyzed by western blot with a rabbit polyclonal antibody to LC3B. Monoclonal antibody against β-ACTIN was used as a loading control. Position of molecular mass markers is indicated on the left. **(F)** Densitometry quantification of LC3-I and LC3-II protein levels from images as those shown in E. Relative levels are expressed as the ratio of LC3-II to LC3-I from images as those shown in E. Bars represent the mean ± standard deviation. Statistical analysis was performed using two-tailed unpaired Student’s t-test (n = 3 n.s. not statistically significant, *p < 0.05, ** p < 0.01, *** p < 0.001)

### The phosphomimetic HERPUD1-S59D mutant promotes ER-lysosomal network with impact in stress cell survival

Because of the increase in the number of functional lysosomes under the expression of HERPUD1-ΔUBL, we investigated if HERPUD1-S59D expressing cells could have a similar output. First, we performed immunofluorescence of LAMP1, observing a significant increase in the number of LAMP1 positive structures in HERPUD1-S59D cells (135.28 ± 21.32), in comparison to HERPUD1-WT (69.03 ± 13.79) and HERPUD1-S59A (58.42 ± 14.55) (Supp. Fig. 9A and 9B). Next, analysis with LysoTracker in HERPUD1-S59D cells (183.35 ± 60.05), in comparison to HERPUD1-WT (118.18 ± 51.00) and HERPUD1-S59A (86.13 ± 41.46) confirmed HERPUD1-S59D cells have a significant increase in the number of lysosomes per cell (Suppl. Fig. 9C and 9D). Similar results were obtained with Magic Red where the quantification showed an increase of almost twice the intensity of Magic Red in HERPUD1-S59D cells (2.25 ± 0.32) in comparison with HERPUD1-WT (1.00 ± 0.15) and HERPUD1-S59A (1.02 ± 0.24) (Suppl. Fig 9E and 9F).

One important aspect of cellular plasticity is the establishment of a network between the ER and the lysosomes that helps in the constant adaptation to particular cellular needs. In this regard, it is known that the ER forms membrane-contact sites (MCSs) with lysosomes (Valm et al. 2017), acting as a potent spatiotemporal organizer of endolysosomal biology (Neefjes, Jongsma, and Berlin 2017). We hypothesized that the expansion of the ER-lysosomal network should be accompanied by the appearance of MCSs between these two organelles. Therefore, we studied the spatial distribution of the ER and lysosomes upon stability of HERPUD1 with the expression of HERPUD1-ΔUBL and HERPUD1-S59D. For this, multiple confocal images were obtained with CALNEXIN and LAMP1 co-staining by immunofluorescence and analyzed in a super-resolution scale. As expected, we observed that similar to HERPUD1-WT expression, expression of HERPUD1-ΔUBL and HERPUD1-S59D form ER-lysosomes MCSs (Fig. 8 middle panel). MCSs are also shown in a three-dimensional (3D) reconstruction with high magnification (Fig. 8 inset lower panel). Altogether, our findings strongly suggest that expansion of ER by HERPUD1 stability not only promotes the remodeling of the lysosomal network but also the formation of MCSs between these two organelles.

**Figure 8.**
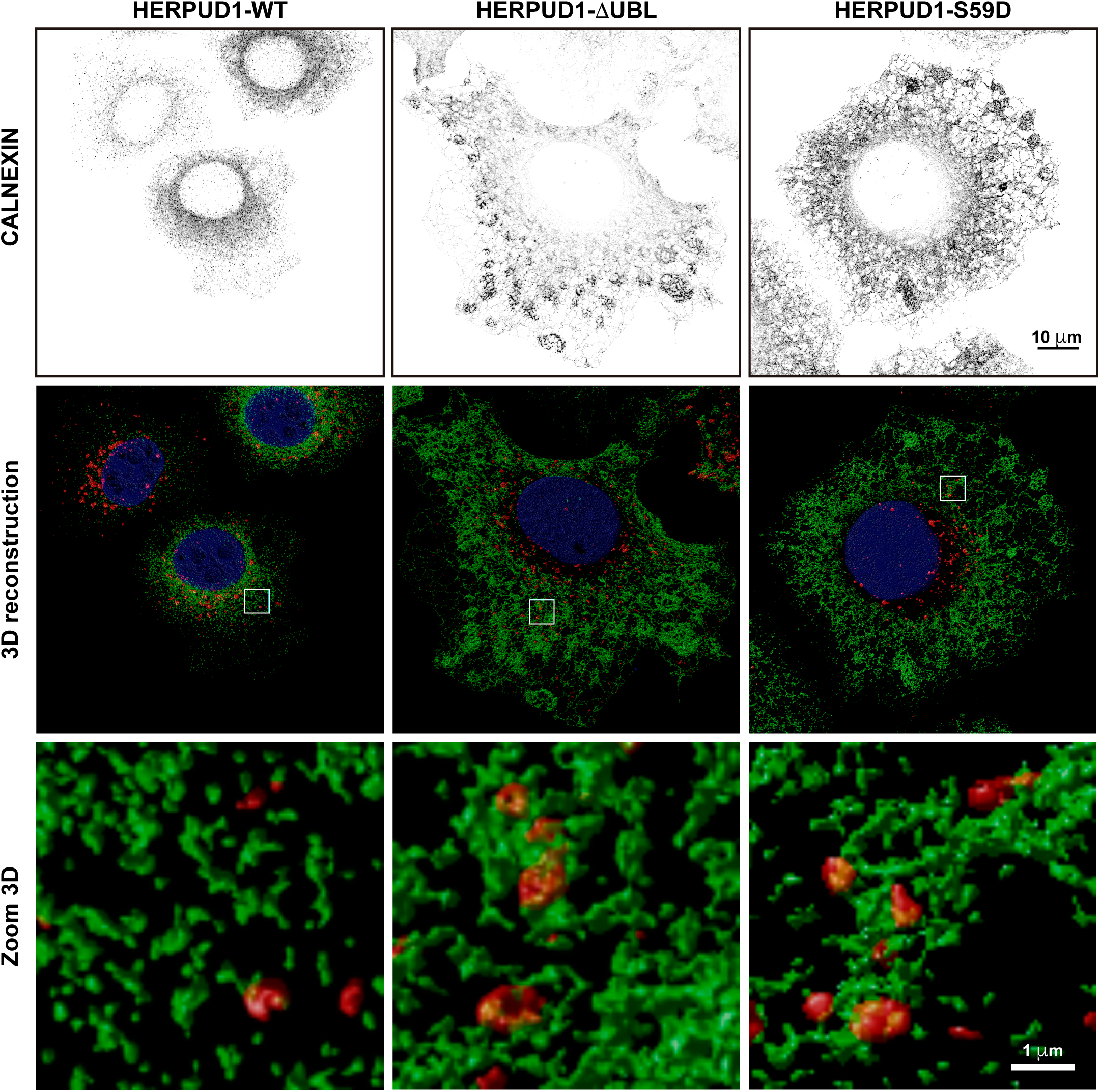
Formation of lysosomal membrane contact sites with the extended ER network in HERPUD1-ΔUBL and HERPUD1-S59D cells. HeLa cells stably expressing HERPUD1-WT-FLAG, HERPUD1-ΔUBL-FLAG or HERPUD1-S59D-FLAG were grown in glass coverslips and then fixed, permeabilized and double-labeled with a rabbit monoclonal antibody against CALNEXIN and a mouse monoclonal antibody against LAMP1 followed by incubation with Alexa-488-conjugated donkey anti-rabbit IgG (green channel), Alexa-594-conjugated donkey anti-mouse IgG (red channel) and DAPI for nuclei staining. All images were acquired using a TCS SP8 laser-scanning confocal microscope in z-series. Upper pannel correspond to inverted z-projection of maximum intensity of CALNEXIN images showing the different ER patterns between cell lines. Middle pannel show 3D reconstructions of CALNEXIN (green channel) and LAMP1 (red channel) images. Scale bar, 10 µm. Lower pannel show a higher magnification of the 3D reconstructions. Scale bar, 1 µm (Note: HERPUD1-WT-FLAG cells are smaller than HERPUD1-S59A-FLAG and HERPUD1-S59D-FLAG).

ER expansion has been known for decades as a trigger for crystalloid ER (Chin et al. 1982) and ER whorls (Feldman, Swarm, and Becker 1981; Schuck et al. 2009), however only recent studies have proposed this phenomenon could be part of a program necessary to overcome ER stress with impact in stress cell survival (Schuck et al. 2009; F. Xu et al. 2021). Moreover, taking in consideration that HERPUD1 stability triggers ER expansion, ER-lysosomal network and MCSs between these two organelles, a protein known to be upregulated under ER stress, we studied the effect of HERPUD1 stability in the cell viability of HeLa cells in response to cisplatin (CDDP). CDDP is the most frequently used chemotherapeutic agent for the treatment of some types of cancers, including cervical cancer in accordance with the model of HeLa cells (Dasari and Bernard Tchounwou 2014). In addition, it is known that alleviation of ER stress attenuates CDDP-induced apoptosis (Wu et al. 2018). Thus, we investigated the resistance to drug-induced cytotoxicity of HERPUD-WT and HERPUD1-S59D cells in response to varying doses of CDDP for 24 h with the Sulforhodamine B (SRB) assay. We found a significant increase in the resistance of HERPUD1-S59D cells in comparison with HERPUD1-WT at 2, 4, 8 and 16 μM of CDDP (Fig. 9A). We corroborated this result measuring apoptosis of these cells in the absence or presence of 10 μM CDDP for 24 h by flow cytometry using Annexin-V staining method. In agreement with the SRB assay, we observed HERPUD1-S59D cells showed less apoptosis compared to HERPUD1-WT cells in response to CDDP treatment (Fig. 9B). These results strongly indicate HERPUD1 stability helps to overcome cisplatin-induced cytotoxicity. In this regard, we propose that this differential response could be mediated by the expansion of the ER/lysosomal network, an aspect that should be investigated in other cancer cell models.

**Figure 9.**
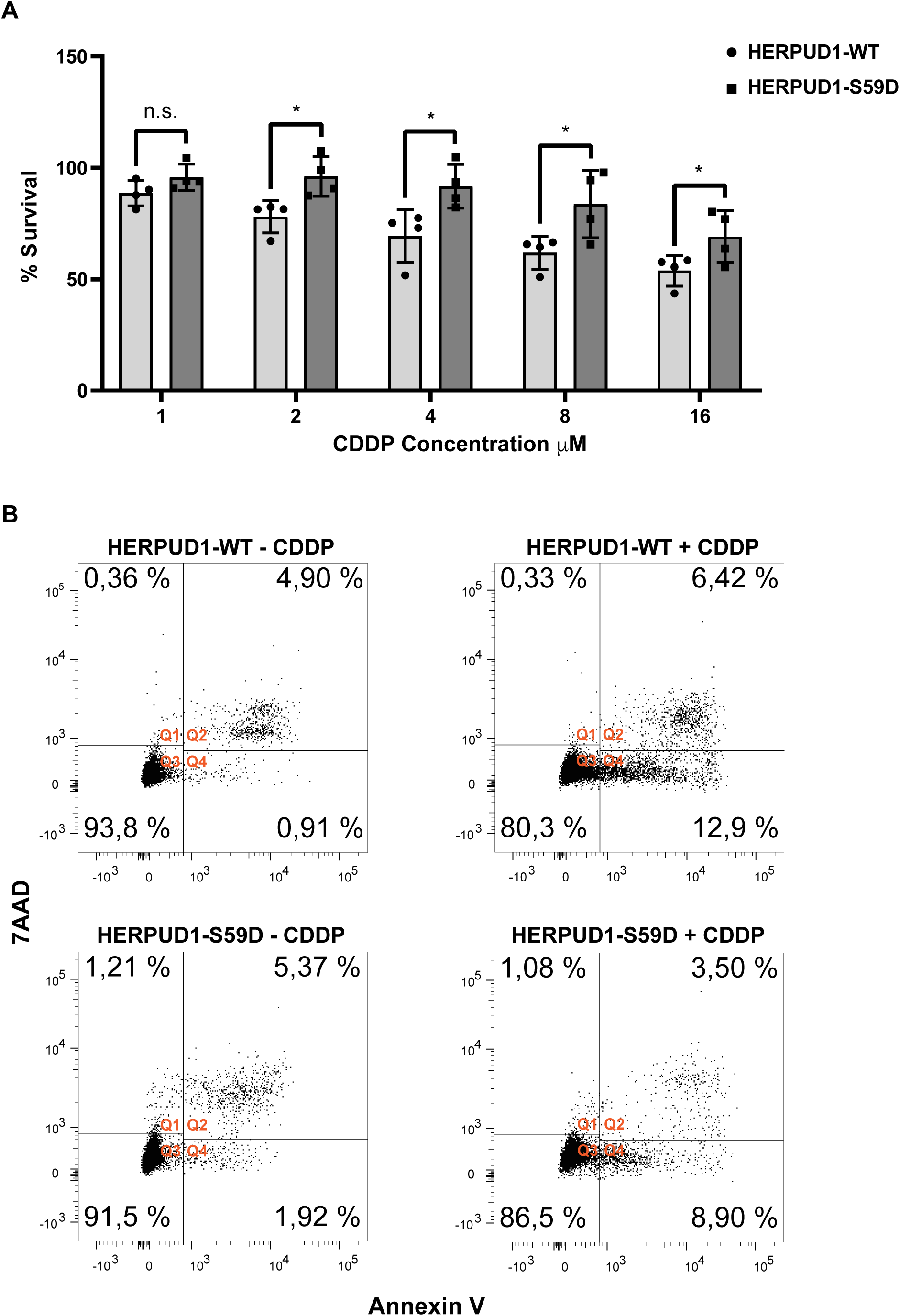
Stabilization of HERPUD1 by S59D mutation decreases cell death mediated by CDDP. **(A)** HeLa cells stably expressing HERPUD1-WT-FLAG or HERPUD1-S59D-FLAG were treated with different concentrations of cisplatin (CDDP) and then a SRB assay was performed. The absorbance values of each point were normalized to control cells (without treatment) and transformed to a percentage. Experiments were performed at least three times, and the results are expressed as mean ± standard deviation. Statistical analysis was performed using two-tailed unpaired Student’s t-test (n = 4 n.s. not statistically significant and *p < 0.05) **(B)** HeLa cells stably expressing HERPUD1-WT-FLAG or HERPUD1-S59D-FLAG were treated or not with CDDP 10 µM for 24 h. Afterwards, cells were collected by trypsinization and stained with Annexin V-Pacific Blue and 7AAD. Scatter plots representing the different quadrants are labeled as Q1 (necrotic cells), Q2 (viable cells), Q3 (dead cells) and Q4 (apoptotic cells). The graph shows the percentage of cells present in each quadrant.

## DISCUSSION

Post-translational responses allow cells to respond swiftly to stress conditions with consequences in their proteome composition. Autophagy and the UPS are two closely related pathways that adjust their functions in response to cellular demands in order to maintain cellular homeostasis (Korolchuk, Menzies, and Rubinsztein 2010). During starvation overall protein degradation rises by activation of both autophagy and the UPS (Mizushima, Yoshimori, and Ohsumi 2011; VerPlank et al. 2019). Indeed, before up-regulation of autophagy, efficient synthesis of new proteins is sustained by degradation of preexisting proteins by the proteasome, which shows that the proteasome also plays a crucial role in cell survival after nutritional stress (Vabulas and Hartl 2005). In agreement with this, it is known that the proteasome activity transits from a latent to an activated state (Asano et al. 2015; Collins and Goldberg 2020). However, if UPS activation upon starvation mediates the degradation of specific proteins that could slow-down autophagy is unknown. Downregulation of HERPUD1 upon nutrient starvation in ATG5-depleted cells opens that alternative. In this regard, it has been previously proposed that HERPUD1 depletion up-regulates autophagy (Quiroga et al. 2013) and the degradation of cytosolic protein aggregates (Miura et al. 2010). These findings support the idea that basal levels of HERPUD1 could act as a break in the activation of basal autophagy.

Under ER stress, HERPUD1 is upregulated even faster than ER chaperones (K. Kokame et al. 2000; Bergmann et al. 2018), where it participates in Endoplasmic Reticulum-Associated Degradation (ERAD) (Schulze et al. 2005). HERPUD1 facilitates the assembly of the HRD1 complex, also known as the retrotranslocon, key in the retrotranslocation of unfolded proteins from the ER to the cytosol for proteasomal degradation upon demand (Leitman et al. 2014; Schulz et al. 2017). However, the function of HERPUD1 in the absence of ER stress is less understood. Here, we propose HERPUD1 acts as a negative regulator of autophagy controlled by its proteasome dependent stability, a mechanism that could operate in the absence of ER stress under the control of phosphorylation of its UBL domain.

One interesting aspect of HERPUD1 is the UBL domain present in its N-terminus region. In general, UBL-containing proteins share the ability to interact with the 19S regulatory particle of the 26S proteasome promoting its activation (Rasmus Hartmann-Petersen and Gordon 2004; Yu and Matouschek 2017; Collins and Goldberg 2020). The UBL domain itself can stimulate multiple proteasome activities in a similar fashion to ubiquitin chains (H. T. Kim and Goldberg 2018; Collins and Goldberg 2020). However, unlike ubiquitin and its homologous (e.g., SUMO and Nedd8), UBL-containing proteins cannot be conjugated to other proteins. The human genome encodes over 60 UBL-containing proteins, where 15 of them have been studied in their ability to bind and regulate proteasome activity (Collins and Goldberg 2020). HERPUD1 is one member of this group, however its role as a positive modulator of the proteasomal activity remains uncharacterized. Moreover, because UBL-containing proteins can stimulate proteasome activity (H. T. Kim and Goldberg 2018; Collins and Goldberg 2020), it would be interesting to explore the effect of HERPUD1-ΔUBL and HERPUD1-S59D on the activity of the proteasome under normal and starvation conditions.

In addition, recent findings show UBL-containing proteins can also play a regulatory role in autophagy, such as USP14 (E. Kim et al. 2018), NUB1 (Guarascio et al. 2020), Elongin B (Antonioli et al. 2016), UHRF1 (Shi et al. 2020), OASL (Batista-Silva et al. 2018), BAT3 (Sebti et al. 2014) and UBQLN (Rothenberg et al. 2010; Şentürk et al. 2019; Yang and Klionsky 2020). Our findings show that the stabilization of HERPUD1 by lacking its UBL domain causes a reduction in LC3-II/LC3-I ratio, which positions HERPUD1 as a member of this growing list of UBL-containing proteins that function as regulators of autophagy. However, how HERPUD1 stability mediates the reduction in LC3-II/LC3-I ratio needs further studies. One aspect to be explored is the finding that HERPUD1 interacts with UBQLN (T. Y. Kim et al. 2008). UBQLN is a cytosolic protein, that in addition to HERPUD1, interacts with LC3 and ubiquitinated cargos in autophagosomes (Rothenberg et al. 2010). Furthermore, it is known that silencing UBQLN leads to a reduction in the lipidation of LC3-I to LC3-II that correlates with a diminished number of autophagosomes (Rothenberg et al. 2010). Because UBQLN binds HERPUD1 independently of its UBL domain (T. Y. Kim et al. 2008), it is possible that an increase in HERPUD1 stability at the ER could mediate the sequestration of UBQLN in this compartment affecting its function in other membranes such as autophagosomes (Rothenberg et al. 2010). In this regard, it has been previously proposed that recruitment of UBQLN to the ER by HERPUD1 could bring the proteasome and the ubiquitinated substrates to specific microdomains of the ER, which could hypothetically be the step that promotes the ERAD pathway (T. Y. Kim et al. 2008). Because our results show that HERPUD1 stability maintain the ER-lysosomal network, additional work is needed to determine if HERPUD1:UBQLN interaction could play a role in the delivery of substrates to the ER-to-lysosomes-associated degradation (ERLAD) (Fregno et al. 2018; Fregno and Molinari 2019), considering the increase in the lysosomal degradation function. As HERPUD1-S59D mimics the effect on HERPUD1 stability we propose that recruitment of UBQLN at the ER is controlled by phosphorylation of HERPUD1.

Ubiquitin and UBL domains are targets of phosphorylation (Kondapalli et al. 2012; Koyano et al. 2014; Wauer et al. 2015). The best-known example is the phosphorylation of the UBL domain of the ubiquitin ligase PARKIN by the Ser/Thr kinase PINK1 on Ser65 (Kondapalli et al. 2012; Koyano et al. 2014). This post-translational modification orchestrates its enzymatic E3 ligase activity (Aguirre et al. 2017), participating in the ubiquitination of mitochondrial proteins during mitophagy. Here, we propose that phosphorylation of the UBL domain in HERPUD1 must affect its noncovalent binding to the proteasome explaining its stability increase, as occurs with HERPUD1-ΔUBL. Further studies are needed to define the physiological triggers of HERPUD1 phosphorylation, and the kinase involved.

This study is the first report indicating HERPUD1 stability mediates ER expansion but not as a consequence of ER stress. In this regard, it is well known that the ER expands to alleviate ER stress (Schuck et al. 2009). However, the ER expansion phenomenon is not always homeostatic. ER expansion requires an adequate supply of membrane lipids although the mechanisms that govern ER biogenesis are yet unclear. For example, the ER expands several folds when B lymphocytes differentiate into antibody-secreting plasma cells (Wiest et al. 1990), when hepatocytes increase its P450 detoxification system (Feldman, Swarm, and Becker 1981), in response to Epidermal Growth Factor (EGF) (Caldieri et al. 2017) and statins (Chin et al. 1982). Interestingly, lipid synthesis activation causes expansion of the ER and resistance to ER stress even in cells lacking the UPR, highlighting the physiological importance of ER membrane biogenesis in homeostasis. A future challenge will be to sort out the connection between HERPUD1 and lipid synthesis, a link recently suggested by genomics (Van Der Laan et al. 2018).

In this regard, an interesting observation to be considered is that HERPUD1 increased stability mimics the effect of statins in reference to the appearance of a crystalloid ER (Chin et al. 1982). Statins are cholesterol reducing agents acting as blockers of the cholesterol biosynthesis by the inhibition of 3-hydroxy-3-methylglutaryl coenzyme A (HMG-CoA) reductase. Interestingly, statins possess beneficial effects in a variety of human diseases. Importantly, a growing number of studies refer to statins as ER stress reducing agents (Mollazadeh et al. 2018; T. Zhang et al. 2018), modulators of autophagy (Ashrafizadeh et al. 2020) and inducers of lysosomal biogenesis (Y. Zhang et al. 2020). However, if statins promote those effects by a mechanism related with HERPUD1 stability is unknown.

Expansion of ER by HERPUD1 stability is correlated with a slow-down in autophagy. In agreement with this, it is known that the knockdown of the two major autophagy regulators, ATG5 and BECN1, likewise trigger ER expansion (Khaminets et al. 2015). Moreover, a similar ER expansion is also observed by ER-phagy inhibition, a selective form of autophagy responsible for the degradation of excess ER (Khaminets et al. 2015; Grumati, Dikic, and Stolz 2018). However, studies investigating if the UPR is activated or not in response to inhibition of autophagosomal biogenesis or ER-Phagy dysfunction are still lacking. While the opposite effect has been reported, observing that excessive ER-Phagy results in activation of the UPR response (Liao et al. 2019).

Our findings also indicate that ER expansion by HERPUD1 stability is accompanied by an increase in the number of functional lysosomes. Because several studies have indicated that UBL-containing proteins cause a positive modulation of the UPS system we can speculate that HERPUD1-ΔUBL and HERPUD1-S59D could have a negative impact in the proteasome activity. Disturbances in the UPS and autophagy function are known to be compensated by lysosome biogenesis (Jackson and Hewitt 2016). In addition, we have already discussed the possibility that HERPUD1 stability could trigger the recruitment of UBQLN to the ER. Interestingly, it has been recently discovered that UBQLN plays a crucial role in the maintenance of the acidic pH of lysosomes and in a closed interplay with the ER (Şentürk et al. 2019; Yang and Klionsky 2020). Further work is needed to determine if HERPUD1:UBQLN interaction could play a role at this level. Furthermore, based on recent findings (Toulmay and Prinz 2011; Henne 2017), it is necessary to explore if MCSs between ER and lysosomes could play a role in the transfer of lipids to lysosomes to compensate for the massive expansion of the ER under HERPUD1 stability by degradation.

Together, our findings highlight novel insights into the possible role of HERPUD1 as a regulator of autophagy and its participation in the maintenance of the ER structure. Moreover, our results suggest that HERPUD1 stabilization promotes the lysosomal function which could promote ER-lysosome intercommunication even in conditions where the UPR is not activated. The mechanism behind this regulation remains to be elucidated, specially the UBL phosphorylation-dependent stabilization of HERPUD1 and its effect on the pathways discussed above. Furthermore, if this regulation has a role in cell pathological conditions it should be analyzed in future studies.

## SUPPLEMENTARY FIGURE LEGENDS

**Supplementary Figure 1.** Table summarizing all proteins identified as significantly differing in abundance (Significance B <0.001) between normal and serum starved conditions for either shLuc or shATG5 treated cells, or both. Number cells are color coded by response to starvation; Blue = reduced abundance, red = increase, white = no change. Number Cells with a black border met significance criteria. Blank number cells correspond to missing data from one or both forward and reverse experiments.

**Supplementary Figure 2. The stabilization of HERPUD1 by its UBL deletion negatively regulates autophagy under starvation**. **(A)** HeLa cells stably expressing HERPUD1-WT-FLAG or HERPUD1-ΔUBL-FLAG untreated (lanes 1 and 2) or treated with EBSS for 4 h in the absence or presence of 100 nM BafA1 (lanes 3 to 6). Detergent-soluble protein extracts were analyzed by western blot with a rabbit polyclonal antibody against LC3. Monoclonal antibody against β-ACTIN (clone BA3R) was used as a loading control. Position of molecular mass markers is indicated on the left. **(B)** Densitometry quantification of LC3-I and LC3-II protein levels from images as those shown in A. Relative levels are expressed as the ratio of LC3-II to LC3-I. Bars represent the mean ± standard deviation. Statistical analysis was performed using two-tailed unpaired Student’s t-test (n = 3 * p < 0.05, ** p < 0.01).

**Supplementary Figure 3. HERPUD1-ΔUBL stabilization alters the ER morphology without changing ER resident protein levels. (A)** HeLa cells stably expressing HERPUD1-WT-FLAG or HERPUD1-ΔUBL-FLAG grown in glass coverslips were fixed, permeabilized and incubated with an antibody against HERPUD1 followed by incubation with Alexa-488-conjugated donkey anti-rabbit IgG. Stained cells were examined by confocal microscopy. Scale bar 10 µm. **(B)** HeLa cells stably expressing HERPUD1-WT-FLAG or HERPUD1-ΔUBL-FLAG were grown in glass bottom culture dishes and labeled with ER-Tracker™ Blue-White DPX. Stained cells were examined by fluorescence microscopy. Images were acquired using a TCS SP8 laser-scanning confocal microscope. Scale bar 10 µm. **(C)** HeLa cells HERPUD1-WT-FLAG or HERPUD1-ΔUBL-FLAG. Detergent-soluble protein extracts were analyzed by western blot with a rabbit monoclonal antibody against CALNEXIN, rat monoclonal antibody against GRP94 and rabbit monoclonal antibody against HERPUD1. Monoclonal antibody against β-ACTIN (clone BA3R) was used as a loading control. Position of molecular mass markers is indicated on the left. **(D)** Densitometry quantification of CALNEXIN and GRP94 protein levels from images as those shown in C. Bars represent the mean ± standard deviation. Statistical analysis was performed using two-tailed unpaired Student’s t-test (n = 3, n.s. not statistically significant).

**Supplementary Figure 4. The stability of HERPUD1 by the deletion of its UBL domain does not trigger endoplasmic reticulum stress.** HeLa WT cells untreated or treated with 2 µM Thapsigargin (Tg) or 5 µg/mL Tunicamycin (Tun) for 4 h (lane 1 to 3) and untreated HeLa cells stably expressing HERPUD1-WT-FLAG or HERPUD1-ΔUBL-FLAG (lane 4 and 5) were lysed. Detergent-soluble protein extracts were analyzed by western blot with a rabbit polyclonal antibody against P-PERK/PERK, rabbit monoclonal antibody against ATF4 and rabbit monoclonal antibody against HERPUD1. Monoclonal antibody against β-ACTIN (clone BA3R) was used as a loading control. Position of molecular mass markers is indicated on the left.

**Supplementary Figure 5. Expression of HERPUD1-ΔUBL increases lysosomal organelle number. (A)** HeLa cells stably expressing HERPUD1-WT-FLAG or HERPUD1-ΔUBL-FLAG grown in glass coverslips were fixed, permeabilized, and double-labeled with mouse monoclonal antibody against LAMP1 and with a goat polyclonal antibody against CATHEPSIN-D (CAT-D) followed by incubation with Alexa-488-conjugated donkey anti-goat IgG (green channel), Alexa-594-conjugated donkey anti-mouse IgG (red channel) and DAPI for nuclei staining. Images were acquired using a TCS SP8 laser-scanning confocal microscope. The third image on each row is the merge of red and green channels; yellow indicates overlapping localization of the red and green channels. Scale bar, 10 µm. **(B)** Quantification of CATHEPSIN-D positive structures per cell. In the scatter plot open circles and open squares represent the average of fluorescence signal for each cell in a frame of HeLa HERPUD1-WT-FLAG (n=559) or HERPUD1-ΔUBL-FLAG (n=700) respectively. Quantified cells were from 3 independent experiments. Statistical analysis was performed using two-tailed unpaired Student’s t-test (*** p < 0.001).

**Supplementary Figure 6. Predicted phosphorylation sites of the UBL domain in HERPUD1**. **(A)** The Ubiquitin-like Domain of HERPUD1 (PDB 1WGD, green color) and ubiquitin (PDB 2MSG, purple color) resolved structures were aligned by PyMOL molecular graphics system. **(B)** Predicted phosphorylated serines are shown in red in the resolved structure of the Ubiquitin-like Domain of HERPUD1 (PDB 1WGD, green color). Each arrow points to the specific residue shown in the figure.

**Supplementary Figure 7. Stabilization of HERPUD1 by S59D mutation decreases autophagosomes.** HeLa cells stably expressing HERPUD1-WT-FLAG or HERPUD1-S59D-FLAG grown in glass coverslips were treated or not with 100 nM BafA1 for 4 h. Cells were fixed, permeabilized and incubated with a polyclonal rabbit antibody against LC3 followed by incubation with Alexa-488-conjugated donkey anti-rabbit IgG. Stained cells were examined by confocal microscopy. Images were acquired using a TCS SP8 laser-scanning confocal microscope. Scale bar 10 µm.

**Supplementary Figure 8. Expression of HERPUD1-S59D increases nuclear and cell size. (A)** Cell size analysis of HeLa cells stably expressing HERPUD1-WT-FLAG, HERPUD1-ΔUBL-FLAG, HERPUD1-S59A-FLAG and HERPUD1-S59D-FLAG. Cells grown in glass coverslips were marked with Wheat Germ Agglutinin (WGA), fixed and stained with DAPI for nuclei visualization. Stained cells were examined by fluorescence microscopy. Cell area was measured by selection of cell contour. **(B)** Analysis of the nuclei size of HeLa HERPUD1-WT-FLAG, HERPUD1-ΔUBL-FLAG, HERPUD1-S59A-FLAG and HERPUD1-S59D-FLAG. Cells grown in glass coverslips were stained with Hoechst for nuclei visualization. Stained cells were examined by fluorescence microscopy. Nuclei area was measured by selection of nuclei contour.

**Supplementary Figure 9. Expression of HERPUD1-S59D increases lysosomal number and function. (A)** HeLa cells stably expressing HERPUD1-WT-FLAG, HERPUD1-S59A-FLAG or HERPUD1-S59D-FLAG grown in glass coverslips were fixed, permeabilized and labeled with a mouse monoclonal antibody against LAMP1 followed by incubation with Alexa-594-conjugated donkey anti-mouse IgG and DAPI for nuclei staining. Stained cells were examined by fluorescence microscopy. Images were acquired using a TCS SP8 laser-scanning confocal microscope. Scale bar 10 µm. **(B)** Quantification of LAMP1 positive structures per cell. In the scatter plot each circle, triangle and rhombus represent the average of positive LAMP1 dots per cell from a frame of HeLa HERPUD1-WT-FLAG (n=559), HERPUD1-S59A-FLAG (n=557) and HERPUD1-S59D-FLAG (n=575) respectively. The quantified cells were from 3 independent experiments. **(C and E)** HeLa cells stably expressing HERPUD1-WT-FLAG, HERPUD1-S59A-FLAG or HERPUD1-S59D-FLAG were grown in glass bottom culture dishes and labeled with **(C)** LysoTracker™ Red DND-99 or **(E)** Magic Red®. For live cell imaging, cell culture medium was replaced with phenol red-free DMEM supplemented with HEPES (10 mM, pH 7.4), and images were acquired with TCS SP8 laser-scanning confocal microscope at 37°C. Scale bar 10 µm**. (D)** Quantification of LysoTracker™ dots per cell. Open circles, triangles and rhombuses represent the average of fluorescence signals for each cell of HeLa HERPUD1-WT-FLAG (n=223), HERPUD1-S59A-FLAG (n=233) or HERPUD1-S59D-FLAG (n=124) respectively. **(F)** Quantification of Magic Red® positive structures. In the scatter plot each open circle, triangle and rhombus represent the average of positive dots for Magic Red per cell from a frame of HeLa HERPUD1-WT-FLAG (n=718), HERPUD1-S59A-FLAG (n=1183) or HERPUD1-S59D-FLAG (n=670) respectively. The quantified cells were from 3 independent experiments. Statistical analysis was performed using two-tailed unpaired Student’s t-test (*** p < 0.001).

## REFERENCES

Aguirre, Jacob D., Karen M. Dunkerley, Pascal Mercier, and Gary S. Shaw. 2017. “Structure of Phosphorylated UBL Domain and Insights into PINK1-Orchestrated Parkin Activation.” Proceedings of the National Academy of Sciences of the United States of America 114(2): 298–303. https://pubmed.ncbi.nlm.nih.gov/28007983/ (March 30, 2021).

Anderson, R. G.W., L. Orci, and M. S. Brown. 1983. “Ultrastructural Analysis of Crystalloid Endoplasmic Reticulum in UT-1 Cells and Its Disappearance in Response to Cholesterol.” Journal of Cell Science.

Antonioli, Manuela, Federica Albiero, Mauro Piacentini, and Gian Maria Fimia. 2016. “Temporal Regulation of Autophagy Response by the CULLIN 4-AMBRA1-CULLIN 5 Axis.” Molecular and Cellular Oncology 3(5). https://pubmed.ncbi.nlm.nih.gov/27857967/ (March 30, 2021).

Asano, Shoh et al. 2015. “A Molecular Census of 26S Proteasomes in Intact Neurons.” Science 347(6220): 439–42. https://pubmed.ncbi.nlm.nih.gov/25613890/ (March 30, 2021).

Ashrafizadeh, Milad, Zahra Ahmadi, Tahereh Farkhondeh, and Saeed Samarghandian. 2020. “Modulatory Effects of Statins on the Autophagy: A Therapeutic Perspective.” Journal of Cellular Physiology 235(4): 3157–68. https://pubmed.ncbi.nlm.nih.gov/31578730/ (March 30, 2021).

Batista-Silva, Leonardo Ribeiro, Rychelle Clayde Affonso Medeiros, Flávio Alves Lara, and Milton Ozório Moraes. 2018. “Type I Interferons, Autophagy and Host Metabolism in Leprosy.” Frontiers in Immunology 9(APR). https://pubmed.ncbi.nlm.nih.gov/29755459/ (March 30, 2021).

Benes, Petr, Vaclav Vetvicka, and Martin Fusek. 2008. “Cathepsin D-Many Functions of One Aspartic Protease.” Critical Reviews in Oncology/Hematology 68(1): 12–28.

Bergmann, Timothy J. et al. 2018. “Chemical Stresses Fail to Mimic the Unfolded Protein Response Resulting from Luminal Load with Unfolded Polypeptides.” Journal of Biological Chemistry 293(15): 5600–5612. https://pubmed.ncbi.nlm.nih.gov/29453283/ (June 21, 2020).

Bertolotti, Anne et al. 2000. “Dynamic Interaction of BiP and ER Stress Transducers in the Unfolded-Protein Response.” Nature Cell Biology 2(6): 326–32. https://pubmed.ncbi.nlm.nih.gov/10854322/ (March 31, 2021).

Blois, Joseph, Adam Smith, and Lee Josephson. 2011. “The Slow Cell Death Response When Screening Chemotherapeutic Agents.” Cancer Chemotherapy and Pharmacology 68(3): 795–803. https://pubmed.ncbi.nlm.nih.gov/21193989/ (March 30, 2021).

Boonacker, Emil et al. 2003. “Rapid Assay to Detect Possible Natural Substrates of Proteases in Living Cells.” BioTechniques 35(4): 766–74. https://pubmed.ncbi.nlm.nih.gov/14579742/ (March 31, 2021).

Borgese, Nica, Maura Francolini, and Erik Snapp. 2006. “Endoplasmic Reticulum Architecture : Structures in Flux.” Current Opinion in Cell Biology 18(4): 358–64. https://pubmed.ncbi.nlm.nih.gov/16806883/ (March 30, 2021).

Bustamante, Hianara A. et al. 2018. “Interplay between the Autophagy-Lysosomal Pathway and the Ubiquitin-Proteasome System: A Target for Therapeutic Development in Alzheimer’s Disease.” Frontiers in Cellular Neuroscience 12. https://pubmed.ncbi.nlm.nih.gov/29867359/ (March 30, 2021).

Bustamante, Hianara A et al. 2020. “The Proteasomal Deubiquitinating Enzyme PSMD14 Regulates Macroautophagy by Controlling Golgi-to-ER Retrograde Transport.” Cells 9(3): 777. https://pubmed.ncbi.nlm.nih.gov/32210007/ (June 23, 2020).

Caldieri, Giusi et al. 2017. “Reticulon 3-Dependent ER-PM Contact Sites Control EGFR Nonclathrin Endocytosis.” Science 356(6338): 617–24. https://pubmed.ncbi.nlm.nih.gov/28495747/ (March 30, 2021).

Camuzard, Olivier, Sabine Santucci-Darmanin, Georges F. Carle, and Valérie Pierrefite-Carle. 2020. “Autophagy in the Crosstalk between Tumor and Microenvironment.” Cancer Letters 490: 143–53. https://pubmed.ncbi.nlm.nih.gov/32634449/ (March 30, 2021).

Cavieres, Viviana A. et al. 2020. “Human Golgi Phosphoprotein 3 Is an Effector of RAB1A and RAB1B.” PLoS ONE 15(8 August). https://pubmed.ncbi.nlm.nih.gov/32790781/ (March 30, 2021).

Chin, D. J. et al. 1982. “Appearance of Crystalloid Endoplasmic Reticulum in Compactin-Resistant Chinese Hamster Cells with a 500-Fold Increase in 3-Hydroxy-3-Methylglutaryl-Coenzyme A Reductase.” Proceedings of the National Academy of Sciences of the United States of America 79(4 I): 1185–89. https://pubmed.ncbi.nlm.nih.gov/6951166/ (March 30, 2021).

Chou, Kai Ming, A. Paul Krapcho, and Miles P. Hacker. 2001. “Impact of the Basic Amine on the Biological Activity and Intracellular Distribution of an Aza-Anthrapyrazole: BBR 3422.” Biochemical Pharmacology 62(10): 1337–43. https://pubmed.ncbi.nlm.nih.gov/11709193/ (March 30, 2021).

Collins, Galen A., and Alfred L. Goldberg. 2020. “Proteins Containing Ubiquitin-like (Ubl) Domains Not Only Bind to 26S Proteasomes but Also Induce Their Activation.” Proceedings of the National Academy of Sciences of the United States of America 117(9): 4664–74. https://pubmed.ncbi.nlm.nih.gov/32071216/ (March 30, 2021).

Dasari, Shaloam, and Paul Bernard Tchounwou. 2014. “Cisplatin in Cancer Therapy: Molecular Mechanisms of Action.” European Journal of Pharmacology 740: 364–78. https://pubmed.ncbi.nlm.nih.gov/25058905/ (March 30, 2021).

Demishtein, Alik et al. 2017. “SQSTM1/P62-Mediated Autophagy Compensates for Loss of Proteasome Polyubiquitin Recruiting Capacity.” Autophagy 13(10): 1697–1708. https://pubmed.ncbi.nlm.nih.gov/28792301/ (March 30, 2021).

De Duve, Christian et al. 1974. “Lysosomotropic Agents.” Biochemical Pharmacology 23(18). https://pubmed.ncbi.nlm.nih.gov/4606365/ (March 30, 2021).

Feldman, Dorothy, Richard L. Swarm, and Janet Becker. 1981. “Ultrastructural Study of Rat Liver and Liver Neoplasms after Long-Term Treatment with Phenobarbital.” Cancer Research.

Fregno, Ilaria et al. 2018. “ ER -to-lysosome-associated Degradation of Proteasome-resistant ATZ Polymers Occurs via Receptor-mediated Vesicular Transport .” The EMBO Journal 37(17). https://pubmed.ncbi.nlm.nih.gov/30076131/ (March 30, 2021).

Fregno, Ilaria, and Maurizio Molinari. 2019. “Proteasomal and Lysosomal Clearance of Faulty Secretory Proteins: ER-Associated Degradation (ERAD) and ER-to-Lysosome-Associated Degradation (ERLAD) Pathways.” Critical Reviews in Biochemistry and Molecular Biology 54(2): 153–63. https://pubmed.ncbi.nlm.nih.gov/31084437/ (March 30, 2021).

Friedman, Jonathan R. et al. 2013. “Endoplasmic Reticulum-Endosome Contact Increases as Endosomes Traffic and Mature.” Molecular Biology of the Cell 24(7): 1030–40. https://pubmed.ncbi.nlm.nih.gov/23389631/ (March 30, 2021).

Golebiowski, Filip et al. 2009. “System-Wide Changes to Sumo Modifications in Response to Heat Shock.” Science Signaling 2(72). https://pubmed.ncbi.nlm.nih.gov/19471022/ (March 30, 2021).

González, Alexis E. et al. 2017. “Autophagosomes Cooperate in the Degradation of Intracellular C-Terminal Fragments of the Amyloid Precursor Protein via the MVB/Lysosomal Pathway.” FASEB Journal 31(6): 2446–59. https://pubmed.ncbi.nlm.nih.gov/28254759/ (June 23, 2020).

Griffiths, Gareth et al. 1988. “The Mannose 6-Phosphate Receptor and the Biogenesis of Lysosomes.” Cell 52(3): 329–41. https://pubmed.ncbi.nlm.nih.gov/2964276/ (March 30, 2021).

Grumati, Paolo, Ivan Dikic, and Alexandra Stolz. 2018. “ER-Phagy at a Glance.” Journal of Cell Science 131(17). https://pubmed.ncbi.nlm.nih.gov/30177506/ (March 30, 2021).

Guarascio, Rosellina et al. 2020. “Negative Regulator of Ubiquitin-Like Protein 1 Modulates the Autophagy-Lysosomal Pathway via P62 to Facilitate the Extracellular Release of Tau Following Proteasome Impairment.” Human Molecular Genetics 29(1): 80–96. https://pubmed.ncbi.nlm.nih.gov/31691796/ (March 30, 2021).

Harding, Heather P., Yuhong Zhang, et al. 2000. “Perk Is Essential for Translational Regulation and Cell Survival during the Unfolded Protein Response.” Molecular Cell 5(5): 897–904. https://pubmed.ncbi.nlm.nih.gov/10882126/ (March 30, 2021).

Harding, Heather P., Isabel Novoa, et al. 2000. “Regulated Translation Initiation Controls Stress-Induced Gene Expression in Mammalian Cells.” Molecular Cell 6(5): 1099–1108. https://pubmed.ncbi.nlm.nih.gov/11106749/ (March 30, 2021).

Hartmann-Petersen, R., and C. Gordon. 2004. “Ubiquitin-Proteasome System.” Cellular and Molecular Life Sciences 61(13). https://pubmed.ncbi.nlm.nih.gov/15224183/ (March 30, 2021).

Hartmann-Petersen, Rasmus, and Colin Gordon. 2004. “Integral UBL Domain Proteins: A Family of Proteasome Interacting Proteins.” Seminars in Cell and Developmental Biology 15(2): 247–59. https://pubmed.ncbi.nlm.nih.gov/15209385/ (March 30, 2021).

Heifetz, Aaron, Roy W. Keenan, and Alan D. Elbein. 1979. “Mechanism of Action of Tunicamycin on the UDP-GlcNAc:Dolichyl-Phosphate GlcNAc-1 -Phosphate Transferase.” Biochemistry 18(11): 2186–92. https://pubs.acs.org/doi/abs/10.1021/bi00578a008 (April 27, 2021).

Henne, William Mike. 2017. “Discovery and Roles of ER-Endolysosomal Contact Sites in Disease.” Advances in experimental medicine and biology 997: 135–47. https://pubmed.ncbi.nlm.nih.gov/28815527/ (March 30, 2021).

Huber, Michael D., and Larry Gerace. 2007. “The Size-Wise Nucleus: Nuclear Volume Control in Eukaryotes.” Journal of Cell Biology 179(4): 583–84. https://pubmed.ncbi.nlm.nih.gov/17998404/ (March 30, 2021).

Jackson, Matthew P., and Eric W. Hewitt. 2016. “Cellular Proteostasis: Degradation of Misfolded Proteins by Lysosomes.” Essays in Biochemistry 60(2): 173–80. https://pubmed.ncbi.nlm.nih.gov/27744333/ (March 30, 2021).

Jia, Rui, and Juan S. Bonifacino. 2019. “Negative Regulation of Autophagy by Uba6-Birc6–Mediated Ubiquitination of Lc3.” eLife 8. https://pubmed.ncbi.nlm.nih.gov/31692446/ (March 30, 2021).

Jin, Seok Min, and Richard J. Youle. 2012. “PINK1-and Parkin-Mediated Mitophagy at a Glance.” Journal of Cell Science 125(4): 795–99. https://pubmed.ncbi.nlm.nih.gov/22448035/ (March 30, 2021).

Jung, Hye Seung et al. 2008. “Loss of Autophagy Diminishes Pancreatic β Cell Mass and Function with Resultant Hyperglycemia.” Cell Metabolism 8(4): 318–24. https://pubmed.ncbi.nlm.nih.gov/18840362/ (April 26, 2021).

Khaminets, Aliaksandr et al. 2015. “Regulation of Endoplasmic Reticulum Turnover by Selective Autophagy.” Nature 522(7556): 354–58. https://pubmed.ncbi.nlm.nih.gov/26040720/ (March 30, 2021).

Khaminets, Aliaksandr, Christian Behl, and Ivan Dikic. 2016. “Ubiquitin-Dependent And Independent Signals In Selective Autophagy.” Trends in Cell Biology 26(1): 6–16. https://pubmed.ncbi.nlm.nih.gov/26437584/ (March 30, 2021).

Kim, Eunkyoung et al. 2018. “Dual Function of USP14 Deubiquitinase in Cellular Proteasomal Activity and Autophagic Flux.” Cell Reports 24(3): 732–43. https://pubmed.ncbi.nlm.nih.gov/30021169/ (March 30, 2021).

Kim, Hyoung Tae, and Alfred L. Goldberg. 2018. “UBL Domain of Usp14 and Other Proteins Stimulates Proteasome Activities and Protein Degradation in Cells.” Proceedings of the National Academy of Sciences of the United States of America 115(50): E11642–50. https://pubmed.ncbi.nlm.nih.gov/30487212/ (March 30, 2021).

Kim, Tae Yeon, Eunmin Kim, Sungjoo Kim Yoon, and Jong Bok Yoon. 2008. “Herp Enhances ER-Associated Protein Degradation by Recruiting Ubiquilins.” Biochemical and Biophysical Research Communications 369(2): 741–46. https://pubmed.ncbi.nlm.nih.gov/18307982/ (March 30, 2021).

Kokame, K., K. L. Agarwal, H. Kato, and T. Miyata. 2000. “Herp, a New Ubiquitin-like Membrane Protein Induced by Endoplasmic Reticulum Stress.” Journal of Biological Chemistry 275(42): 32846–53. https://pubmed.ncbi.nlm.nih.gov/10922362/ (June 21, 2020).

Kokame, Koichi, Hisao Kato, and Toshiyuki Miyata. 2001. “Identification of ERSE-II, a New Cis-Acting Element Responsible for the ATF6-Dependent Mammalian Unfolded Protein Response.” Journal of Biological Chemistry 276(12): 9199–9205. https://pubmed.ncbi.nlm.nih.gov/11112790/ (March 30, 2021).

Kondapalli, Chandana et al. 2012. “PINK1 Is Activated by Mitochondrial Membrane Potential Depolarization and Stimulates Parkin E3 Ligase Activity by Phosphorylating Serine 65.” Open Biology 2(MAY). https://pubmed.ncbi.nlm.nih.gov/22724072/ (March 30, 2021).

Korolchuk, Viktor I., Fiona M. Menzies, and David C. Rubinsztein. 2010. “Mechanisms of Cross-Talk between the Ubiquitin-Proteasome and Autophagy-Lysosome Systems.” FEBS Letters 584(7): 1393–98. https://pubmed.ncbi.nlm.nih.gov/20040365/ (March 30, 2021).

Koyano, Fumika et al. 2014. “Ubiquitin Is Phosphorylated by PINK1 to Activate Parkin.” Nature 510(7503): 162–66. https://pubmed.ncbi.nlm.nih.gov/24784582/ (June 24, 2020).

Van Der Laan, Sander W. et al. 2018. “From Lipid Locus to Drug Target through Human Genomics.” Cardiovascular Research 114(9): 1258–70. https://pubmed.ncbi.nlm.nih.gov/29800275/ (March 30, 2021).

Lee, Crystal A., and Craig Blackstone. 2020. “ER Morphology and Endo-Lysosomal Crosstalk: Functions and Disease Implications.” Biochimica et Biophysica Acta - Molecular and Cell Biology of Lipids 1865(1). https://pubmed.ncbi.nlm.nih.gov/31678515/ (March 30, 2021).

Lee, Do Hee. 1998. “Proteasome Inhibitors: Valuable New Tools for Cell Biologists.” Trends in Cell Biology 8(10): 397–403. https://pubmed.ncbi.nlm.nih.gov/9789328/ (March 30, 2021).

Leitman, Julia et al. 2014. “Herp Coordinates Compartmentalization and Recruitment of HRD1 and Misfolded Proteins for ERAD.” Molecular Biology of the Cell 25(7): 1050–60. https://pubmed.ncbi.nlm.nih.gov/24478453/ (June 23, 2020).

Liao, Yangjie et al. 2019. “Excessive ER-Phagy Mediated by the Autophagy Receptor FAM134B Results in ER Stress, the Unfolded Protein Response, and Cell Death in HeLa Cells.” Journal of Biological Chemistry 294(52): 20009–23. https://pubmed.ncbi.nlm.nih.gov/31748416/ (March 30, 2021).

Marzec, Michal, Davide Eletto, and Yair Argon. 2012. “GRP94: An HSP90-like Protein Specialized for Protein Folding and Quality Control in the Endoplasmic Reticulum.” Biochimica et Biophysica Acta - Molecular Cell Research 1823(3): 774–87. https://pubmed.ncbi.nlm.nih.gov/22079671/ (March 31, 2021).

Miettinen, Teemu P., and Mikael Björklund. 2015. “Mevalonate Pathway Regulates Cell Size Homeostasis and Proteostasis through Autophagy.” Cell Reports 13(11): 2610–20. https://pubmed.ncbi.nlm.nih.gov/26686643/ (March 30, 2021).

Miura, Hikari et al. 2010. “Deletion of Herp Facilitates Degradation of Cytosolic Proteins.” Genes to Cells 15(8): 843–53. https://pubmed.ncbi.nlm.nih.gov/20604806/ (March 30, 2021).

Mizushima, Noboru, Beth Levine, Ana Maria Cuervo, and Daniel J. Klionsky. 2008. “Autophagy Fights Disease through Cellular Self-Digestion.” Nature 451(7182): 1069–75. https://pubmed.ncbi.nlm.nih.gov/18305538/ (March 30, 2021).

Mizushima, Noboru, Tamotsu Yoshimori, and Yoshinori Ohsumi. 2011. “The Role of Atg Proteins in Autophagosome Formation.” Annual Review of Cell and Developmental Biology 27: 107–32. https://pubmed.ncbi.nlm.nih.gov/21801009/ (March 30, 2021).

Mollazadeh, Hamid et al. 2018. “The Effect of Statin Therapy on Endoplasmic Reticulum Stress.” Pharmacological Research 137: 150–58. https://pubmed.ncbi.nlm.nih.gov/30312664/ (March 30, 2021).

Moreau, Kevin, Shouqing Luo, and David C. Rubinsztein. 2010. “Cytoprotective Roles for Autophagy.” Current Opinion in Cell Biology 22(2): 206–11. https://pubmed.ncbi.nlm.nih.gov/20045304/ (March 30, 2021).

Murrow, Lyndsay, and Jayanta Debnath. 2013. “Autophagy as a Stress-Response and Quality-Control Mechanism: Implications for Cell Injury and Human Disease.” Annual Review of Pathology: Mechanisms of Disease 8: 105–37. https://pubmed.ncbi.nlm.nih.gov/23072311/ (March 30, 2021).

Neefjes, Jacques, Marlieke M.L. Jongsma, and Ilana Berlin. 2017. “Stop or Go? Endosome Positioning in the Establishment of Compartment Architecture, Dynamics, and Function.” Trends in Cell Biology 27(8): 580–94. https://pubmed.ncbi.nlm.nih.gov/28363667/ (March 30, 2021).

Ou, W. J. et al. 1995. “Conformational Changes Induced in the Endoplasmic Reticulum Luminal Domain of Calnexin by Mg-ATP and Ca2+.” Journal of Biological Chemistry 270(30): 18051–59. https://pubmed.ncbi.nlm.nih.gov/7629114/ (March 30, 2021).

Quiroga, Clara et al. 2013. “Herp Depletion Protects from Protein Aggregation by Up-Regulating Autophagy.” Biochimica et Biophysica Acta - Molecular Cell Research 1833(12): 3295–3305. https://pubmed.ncbi.nlm.nih.gov/24120520/ (March 30, 2021).

Rothenberg, Cara et al. 2010. “Ubiquilin Functions in Autophagy and Is Degraded by Chaperone-Mediated Autophagy.” Human Molecular Genetics 19(16): 3219–32. https://pubmed.ncbi.nlm.nih.gov/20529957/ (March 30, 2021).

Sai, Xiaorei et al. 2003. “The Ubiquitin-like Domain of Herp Is Involved in Herp Degradation, but Not Necessary for Its Enhancement of Amyloid β-Protein Generation.” FEBS Letters 553(1–2): 151–56. https://pubmed.ncbi.nlm.nih.gov/14550564/ (March 30, 2021).

Sauvé, Véronique et al. 2018. “Mechanism of Parkin Activation by Phosphorylation.” Nature Structural and Molecular Biology 25(7): 623–30. https://pubmed.ncbi.nlm.nih.gov/29967542/ (March 30, 2021).

Schindelin, Johannes et al. 2012. “Fiji: An Open-Source Platform for Biological-Image Analysis.” Nature Methods 9(7): 676–82. https://pubmed.ncbi.nlm.nih.gov/22743772/ (March 30, 2021).

Schuck, Sebastian et al. 2009. “Membrane Expansion Alleviates Endoplasmic Reticulum Stress Independently of the Unfolded Protein Response.” Journal of Cell Biology 187(4): 525–36. https://pubmed.ncbi.nlm.nih.gov/19948500/ (March 30, 2021).

Schulz, Jasmin et al. 2017. “Conserved Cytoplasmic Domains Promote Hrd1 Ubiquitin Ligase Complex Formation for ER-Associated Degradation (ERAD).” Journal of Cell Science 130(19): 3322–35. https://pubmed.ncbi.nlm.nih.gov/28827405/ (March 30, 2021).

Schulze, Andrea et al. 2005. “The Ubiquitin-Domain Protein HERP Forms a Complex with Components of the Endoplasmic Reticulum Associated Degradation Pathway.” Journal of Molecular Biology 354(5): 1021–27. https://pubmed.ncbi.nlm.nih.gov/16289116/ (March 30, 2021).

Sebti, Salwa et al. 2014. “BAT3 Modulates P300-Dependent Acetylation of P53 and Autophagy-Related Protein 7 (ATG7) during Autophagy.” Proceedings of the National Academy of Sciences of the United States of America 111(11): 4115–20. https://pubmed.ncbi.nlm.nih.gov/24591579/ (March 30, 2021).

Şentürk, Mümine et al. 2019. “Ubiquilins Regulate Autophagic Flux through MTOR Signalling and Lysosomal Acidification.” Nature Cell Biology 21(3): 384–96. https://pubmed.ncbi.nlm.nih.gov/30804504/ (March 30, 2021).

Shi, Xiaojuan et al. 2020. “Silencing UHRF1 Enhances Cell Autophagy to Prevent Articular Chondrocytes from Apoptosis in Osteoarthritis through PI3K/AKT/MTOR Signaling Pathway.” Biochemical and Biophysical Research Communications 529(4): 1018–24. https://pubmed.ncbi.nlm.nih.gov/32819559/ (March 30, 2021).

Swaney, Danielle L, Ricard A Rodríguez-Mias, and Judit Villén. 2015. “Phosphorylation of Ubiquitin at Ser65 Affects Its Polymerization, Targets, and Proteome-wide Turnover.” EMBO reports 16(9): 1131–44. https://pubmed.ncbi.nlm.nih.gov/26142280/ (June 24, 2020).

Tanida, Isei, Takashi Ueno, and Eiki Kominami. 2008. “LC3 and Autophagy.” Methods in Molecular Biology 445: 77–88. https://pubmed.ncbi.nlm.nih.gov/18425443/ (March 30, 2021).

Tapia, Diego et al. 2019. “KDEL Receptor Regulates Secretion by Lysosome Relocation-and Autophagy-Dependent Modulation of Lipid-Droplet Turnover.” Nature Communications 10(1). https://pubmed.ncbi.nlm.nih.gov/30760704/ (March 30, 2021).

Thastrup, O et al. 1990. “Thapsigargin, a Tumor Promoter, Discharges Intracellular Ca2+ Stores by Specific Inhibition of the Endoplasmic Reticulum Ca2(+)-ATPase.” Proceedings of the National Academy of Sciences 87(7).

Thayer, Julia A. et al. 2020. “The PARK10 Gene USP24 Is a Negative Regulator of Autophagy and ULK1 Protein Stability.” Autophagy 16(1): 140–53. https://pubmed.ncbi.nlm.nih.gov/30957634/ (March 30, 2021).

Toulmay, Alexandre, and William A. Prinz. 2011. “Lipid Transfer and Signaling at Organelle Contact Sites: The Tip of the Iceberg.” Current Opinion in Cell Biology 23(4): 458–63. https://pubmed.ncbi.nlm.nih.gov/21555211/ (March 30, 2021).

Tsubuki, Satoshi et al. 1996. “Differential Inhibition of Calpain and Proteasome Activities by Peptidyl Aldehydes of Di-Leucine and Tri-Leucine.” Journal of Biochemistry 119(3): 572–76. https://pubmed.ncbi.nlm.nih.gov/8830056/ (March 30, 2021).

Vabulas, Ramunas M., and F. Ulrich Hartl. 2005. “Cell Biology: Protein Synthesis upon Acute Nutrient Restriction Relies on Proteasome Function.” Science 310(5756): 1960–63. https://pubmed.ncbi.nlm.nih.gov/16373576/ (March 30, 2021).

Valm, Alex M. et al. 2017. “Applying Systems-Level Spectral Imaging and Analysis to Reveal the Organelle Interactome.” Nature 546(7656): 162–67. https://pubmed.ncbi.nlm.nih.gov/28538724/ (March 30, 2021).

VerPlank, Jordan J.S., Sudarsanareddy Lokireddy, Jinghui Zhao, and Alfred L. Goldberg. 2019. “26S Proteasomes Are Rapidly Activated by Diverse Hormones and Physiological States That Raise CAMP and Cause Rpn6 Phosphorylation.” Proceedings of the National Academy of Sciences of the United States of America 116(10): 4228–37. https://pubmed.ncbi.nlm.nih.gov/30782827/ (March 30, 2021).

Walczak, Marta, and Sascha Martens. 2013. “Dissecting the Role of the Atg12-Atg5-Atg16 Complex during Autophagosome Formation.” Autophagy 9(3): 424–25. https://pubmed.ncbi.nlm.nih.gov/23321721/ (March 30, 2021).

Wauer, Tobias et al. 2015. “Ubiquitin Ser65 Phosphorylation Affects Ubiquitin Structure, Chain Assembly and Hydrolysis.” The EMBO Journal 34(3): 307–25. https://pubmed.ncbi.nlm.nih.gov/25527291/ (March 30, 2021).

Wiest, D. L. et al. 1990. “Membrane Biogenesis during B Cell Differentiation: Most Endoplasmic Reticulum Proteins Are Expressed Coordinately.” Journal of Cell Biology 110(5): 1501–11. https://pubmed.ncbi.nlm.nih.gov/2335560/ (March 30, 2021).

Wu, Yuping et al. 2018. “Alleviation of Endoplasmic Reticulum Stress Protects against Cisplatin-Induced Ovarian Damage.” Reproductive Biology and Endocrinology 16(1). https://pubmed.ncbi.nlm.nih.gov/30176887/ (March 30, 2021).

Xu, Fang et al. 2021. “COPII Mitigates ER Stress by Promoting Formation of ER Whorls.” Cell Research 31(2): 141–56. https://pubmed.ncbi.nlm.nih.gov/32989223/ (March 30, 2021).

Xu, Haoxing, and Dejian Ren. 2015. “Lysosomal Physiology.” Annual Review of Physiology 77: 57–80. https://pubmed.ncbi.nlm.nih.gov/25668017/ (March 30, 2021).

Yan, Long et al. 2014. “Ube2g2-Gp78-Mediated HERP Polyubiquitylation Is Involved in ER Stress Recovery.” Journal of Cell Science 127(7): 1417–27. https://pubmed.ncbi.nlm.nih.gov/24496447/ (June 21, 2020).

Yang, Ying, and Daniel J. Klionsky. 2020. “A Novel Role of UBQLNs (Ubiquilins) in Regulating Autophagy, MTOR Signaling and v-ATPase Function.” Autophagy 16(1): 1–2. https://pubmed.ncbi.nlm.nih.gov/31516068/ (March 30, 2021).

Yin, Yili et al. 2012. “SUMO-Targeted Ubiquitin E3 Ligase RNF4 Is Required for the Response of Human Cells to DNA Damage.” Genes and Development 26(11): 1196–1208. https://pubmed.ncbi.nlm.nih.gov/22661230/ (March 30, 2021).

Yoshida, Hiderou et al. 2001. “XBP1 MRNA Is Induced by ATF6 and Spliced by IRE1 in Response to ER Stress to Produce a Highly Active Transcription Factor.” Cell 107(7): 881–91. https://pubmed.ncbi.nlm.nih.gov/11779464/ (March 30, 2021).

Yu, Houqing, and Andreas Matouschek. 2017. “Recognition of Client Proteins by the Proteasome.” Annual Review of Biophysics 46: 149–73. https://pubmed.ncbi.nlm.nih.gov/28301771/ (March 30, 2021).

Zhang, Tao et al. 2018. “HMG-CoA Reductase Inhibitors Relieve Endoplasmic Reticulum Stress by Autophagy Inhibition in Rats with Permanent Brain Ischemia.” Frontiers in Neuroscience 12(JUN). https://pubmed.ncbi.nlm.nih.gov/29970982/ (March 30, 2021).

Zhang, Youzhi et al. 2020. “Simvastatin Improves Lysosome Function via Enhancing Lysosome Biogenesis in Endothelial Cells.” Frontiers in Bioscience - Landmark.

Zhu, K., K. Dunner, and D. J. McConkey. 2010. “Proteasome Inhibitors Activate Autophagy as a Cytoprotective Response in Human Prostate Cancer Cells.” Oncogene 29(3): 451–62. https://pubmed.ncbi.nlm.nih.gov/19881538/ (March 30, 2021).

