## Supplemental Data for "Negative modulation of macroautophagy by HERPUD1 is counteracted by an increased ER-lysosomal network with impact in drug-induced stress cell survival"

| Gene names | Log2 Starvation /Norm shLuc | Log2 Starvation /Norm shATG5 | Protein names |
|---|---|---|---|
| HERPUD1 | -2.3755 | -2.9502 | Homocysteine-responsive endoplasmic reticulum-resident ubiquitin-like domain member 1 protein |
| SQSTM1 | -1.9500 | -1.7523 | Sequestosome-1 |
| RRM2 | -1.7683 | -1.4810 | Ribonucleoside-diphosphate reductase subunit M2 |
| CHCHD2;CHCHD2P9 | -1.5471 | -1.2523 | Coiled-coil-helix-coiled-coil-helix domain-containing protein 2, mitochondrial |
| CYP1B1 | -1.5041 | -1.1918 | Cytochrome P450 1B1 |
| JAK1 | -1.3608 | -1.2387 | Tyrosine-protein kinase JAK1 |
| PIK3C3 | -1.2089 | | Phosphatidylinositol 3-kinase |
| FAM127A | | -1.1671 | Protein FAM127A |
| UBE2S | | -1.1280 | Ubiquitin-conjugating enzyme E2 S |
| DSP | -1.1525 | -1.0391 | Desmoplakin |
| HMOX1 | -1.3142 | -0.7418 | Heme oxygenase 1 |
| UCK2 | -1.1414 | -0.9024 | Uridine-cytidine kinase 2 |
| DIDO1 | | -1.0206 | Death-inducer obliterator 1 |
| CCNB1 | | -0.9982 | G2/mitotic-specific cyclin-B1 |
| DNAJA1 | -1.1604 | -0.8325 | DnaJ homolog subfamily A member 1 |
| CTNNA2 | -0.9902 | | Catenin alpha-2 |
| PRSS56 | -0.7467 | -1.2140 | Serine protease 56 |
| DDX5 | -1.0243 | -0.8596 | Probable ATP-dependent RNA helicase DDX5 |
| CDKN2A | -1.0032 | -0.8376 | Cyclin-dependent kinase inhibitor 2A, isoforms 1/2/3 |
| DNAJB4 | -1.0796 | -0.6709 | DnaJ homolog subfamily B member 4 |
| HSPA1A | -1.0232 | -0.6260 | Heat shock 70 kDa protein 1A/1B |
| HNRNPK | -0.7659 | -0.8384 | Heterogeneous nuclear ribonucleoprotein K |
| MAT2A | -0.8386 | -0.7585 | S-adenosylmethionine synthase isoform type-2 |
| NTPCR | -1.3866 | -0.2104 | Cancer-related nucleoside-triphosphatase |
| CTNNA1 | -0.7467 | -0.7750 | Catenin alpha-1 |
| YTHDF2 | -0.8372 | -0.6683 | YTH domain-containing family protein 2 |
| LARP4 | -1.0016 | -0.4818 | La-related protein 4 |
| DFFA | -0.8074 | -0.6549 | DNA fragmentation factor subunit alpha |
| KPNA2 | -0.8162 | -0.6413 | Importin subunit alpha-1 |
| CSDE1 | -0.7522 | -0.6710 | Cold shock domain-containing protein E1 |
| PDLIM5 | -0.7219 | -0.6991 | PDZ and LIM domain protein 5 |
| SLC39A14 | -0.7333 | -0.6765 | Zinc transporter ZIP14 |
| EPPK1 | -0.8574 | -0.5506 | Epiplakin |
| KPNA1 | -0.5970 | -0.8046 | Importin subunit alpha-5 |
| DNAJB1 | -0.8146 | -0.5555 | DnaJ homolog subfamily B member 1 |
| BAG3 | -0.7414 | -0.6209 | BAG family molecular chaperone regulator 3 |
| EHD4 | -0.5058 | -0.8238 | EH domain-containing protein 4 |
| VPS13A | -0.6579 | | Vacuolar protein sorting-associated protein 13A |
| CTTN | -0.7476 | -0.5014 | Src substrate cortactin |
| CKAP5 | -0.7423 | -0.4996 | Cytoskeleton-associated protein 5 |
| INF2 | -0.7301 | -0.5117 | Inverted formin-2 |
| DIABLO | -0.7455 | -0.4922 | Diablo homolog, mitochondrial |
| ZC3HAV1 | -0.6140 | -0.5688 | Zinc finger CCCH-type antiviral protein 1 |
| ASNS | -0.5693 | -0.6099 | Asparagine synthetase [glutamine-hydrolyzing] |
| TSR1 | -0.7308 | -0.4485 | Pre-rRNA-processing protein TSR1 homolog |
| CYR61 | -0.1370 | -1.0238 | Protein CYR61 |
| PAK2 | -0.5284 | -0.6034 | Serine/threonine-protein kinase PAK 2 |
| FXR1 | -0.5265 | -0.6019 | Fragile X mental retardation syndrome-related protein 1 |
| EIF5 | -0.4853 | -0.6181 | Eukaryotic translation initiation factor 5 |
| SSRP1 | -0.7011 | -0.3745 | FACT complex subunit SSRP1 |
| PICALM | -0.3923 | -0.6463 | Phosphatidylinositol-binding clathrin assembly protein |
| RRM1 | -0.7963 | -0.1904 | Ribonucleoside-diphosphate reductase large subunit |
| KRT18 | -0.6322 | -0.3419 | Keratin, type I cytoskeletal 18 |
| TPT1 | -0.6253 | -0.3430 | Translationally-controlled tumor protein |
| EIF4B | -0.4945 | -0.4216 | Eukaryotic translation initiation factor 4B |
| DDX3X;DDX3Y | -0.4678 | -0.3529 | ATP-dependent RNA helicase DDX3X |
| DDX17 | -0.4658 | -0.3408 | Probable ATP-dependent RNA helicase DDX17 |
| WARS | -0.4621 | -0.3410 | Tryptophan--tRNA ligase, cytoplasmic |
| HSPA6 | -0.4815 | -0.3151 | Heat shock 70 kDa protein 6 |
| LMNB1 | -0.4677 | -0.2722 | Lamin-B1 |
| HSPH1 | -0.4016 | -0.2609 | Heat shock protein 105 kDa |
| NONO | -0.3935 | -0.2097 | Non-POU domain-containing octamer-binding protein |
| CENPF | -0.5236 | 0.0339 | Centromere protein F |
| KPRP | 0.7616 | -1.1857 | Keratinocyte proline-rich protein |
| S100A8 | 0.3830 | -0.5917 | Protein S100-A8 |
| PDLIM7 | -0.4601 | 0.6020 | PDZ and LIM domain protein 7 |
| APOBEC3C | -0.6987 | 1.1531 | DNA dC->dU-editing enzyme APOBEC-3C |
| HNRNPA2B1 | 0.4558 | 0.1988 | Heterogeneous nuclear ribonucleoproteins A2/B1 |
| SYPL1 | 0.9425 | -0.2530 | Synaptophysin-like protein 1 |
| NOP56 | 0.8012 | -0.0723 | Nucleolar protein 56 |
| EFTUD2 | 0.4682 | 0.3682 | 116 kDa U5 small nuclear ribonucleoprotein component |
| TCOF1 | 0.5133 | 0.3271 | Treacle protein |
| SNRPA1 | 0.4857 | 0.4099 | U2 small nuclear ribonucleoprotein A |
| BCCIP | -0.0579 | 0.9899 | BRCA2 and CDKN1A-interacting protein |
| DDAH2 | 0.8945 | 0.0728 | N(G),N(G)-dimethylarginine dimethylaminohydrolase 2 |
| SLC4A7 | 0.2193 | 0.7783 | Sodium bicarbonate cotransporter 3 |
| PSIP1 | 0.7405 | 0.3509 | PC4 and SFRS1-interacting protein |
| HIST1H1E | 0.7938 | 0.3070 | Histone H1.4 |
| BRIX1 | 0.7510 | 0.4148 | Ribosome biogenesis protein BRX1 homolog |
| HIST1H1C;HIST1H1D | 0.7096 | 0.4598 | Histone H1.2 |
| FAM96A | -0.0920 | 1.3367 | MIP18 family protein FAM96A |
| MRTO4 | 0.7314 | 0.5427 | mRNA turnover protein 4 homolog |
| GTPBP4 | 1.0446 | 0.2341 | Nucleolar GTP-binding protein 1 |
| HIST1H1B | 0.8032 | 0.5083 | Histone H1.5 |
| CXADR | 1.4132 | -0.0661 | Coxsackievirus and adenovirus receptor |
| MAN2A1 | 0.2853 | 1.1192 | Alpha-mannosidase 2 |
| GYG1 | 0.8479 | 0.5741 | Glycogenin-1 |
| HMGN1 | 1.0140 | 0.4165 | Non-histone chromosomal protein HMG-14 |
| GLI2 | 0.6774 | 0.7593 | Zinc finger protein GLI2 |
| PKP3 | -0.5195 | 1.9642 | Plakophilin-3 |
| DSG1 | | 0.7253 | Desmoglein-1 |
| PDCD4 | 0.6855 | 0.8296 | Programmed cell death protein 4 |
| NOP2 | 1.0216 | 0.5255 | Probable 28S rRNA (cytosine(4447)-C(5))-methyltransferase |
| ALDH1A3 | 1.4997 | 0.0761 | Aldehyde dehydrogenase family 1 member A3 |
| TAF15 | 0.8080 | | TATA-binding protein-associated factor 2N |
| MRPS26 | 1.4780 | 0.1647 | 28S ribosomal protein S26, mitochondrial |
| ERC1 | 0.8614 | | ELKS/Rab6-interacting/CAST family member 1 |
| HMGN3;HMGN2 | 0.9721 | 0.7621 | High mobility group nucleosome-binding domain-containing prot. 3 |
| DDX27 | 1.0440 | 0.7351 | Probable ATP-dependent RNA helicase DDX27 |
| VIM | 0.7521 | 1.1510 | Vimentin |
| RSL1D1 | 1.2941 | 0.7582 | Ribosomal L1 domain-containing protein 1 |
| RALY | 1.2345 | 0.8586 | RNA-binding protein Raly |
| FBL | 1.6435 | 0.8101 | rRNA 2-O-methyltransferase fibrillarin |
| HNRNPC;HNRNPCL1 | 1.5508 | 0.9595 | Heterogeneous nuclear ribonucleoproteins C1/C2 |
| NUMA1 | 1.4976 | 1.0158 | Nuclear mitotic apparatus protein 1 |
| SBSN | 1.3332 | | Suprabasin |
| H2AFV;H2AFZ | 1.7766 | 1.3893 | Histone H2A.V |
| H3F3A | 2.0758 | 1.8649 | Histone H3.3 |
| HIST1H2AC;HIST3H2A;HIST1H2AB | 2.5651 | 2.0782 | Histone H2A type 1-C |
| HIST2H3A | 2.8848 | 1.8520 | Histone H3.2 |
| HIST1H2BL;HIST1H2BM;HIST1H2BN;HIST1H2BH;HIST2H2BF;HIST1H2BC;HIST1H2BD;HIST1H2BK;H2BFS | 2.9014 | 2.2069 | Histone H2B |
| HIST1H4A | 2.8226 | 2.3639 | Histone H4 |
| HIST2H2AC;HIST2H2AA3 | 2.9450 | 2.2908 | Histone H2A type 2-C |
| HIST2H2BE;HIST1H2BB;HIST1H2BO;HIST1H2BJ;HIST3H2BB | 3.0875 | 2.3765 | Histone H2B type 2-E |
| HIST2H3PS2 | 2.8909 | 3.0398 | Histone H3 |
| H2AFY | 3.1737 | 2.8956 | Core histone macro-H2A.1 |
| HIST1H3A;HIST3H3;H3F3A;H3F3C | 3.3200 | 2.7767 | Histone H3.1 |

**Supplementary Figure 1**

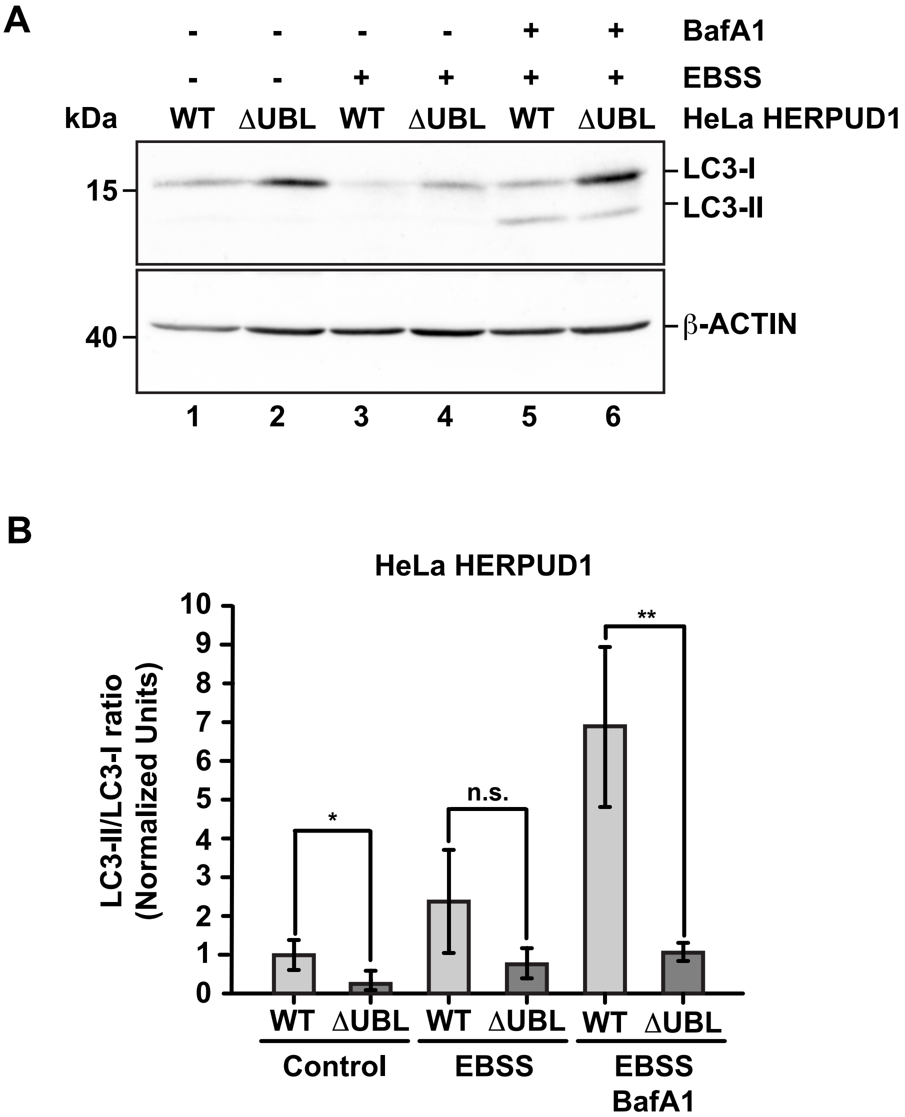


Supplementary Figure 2

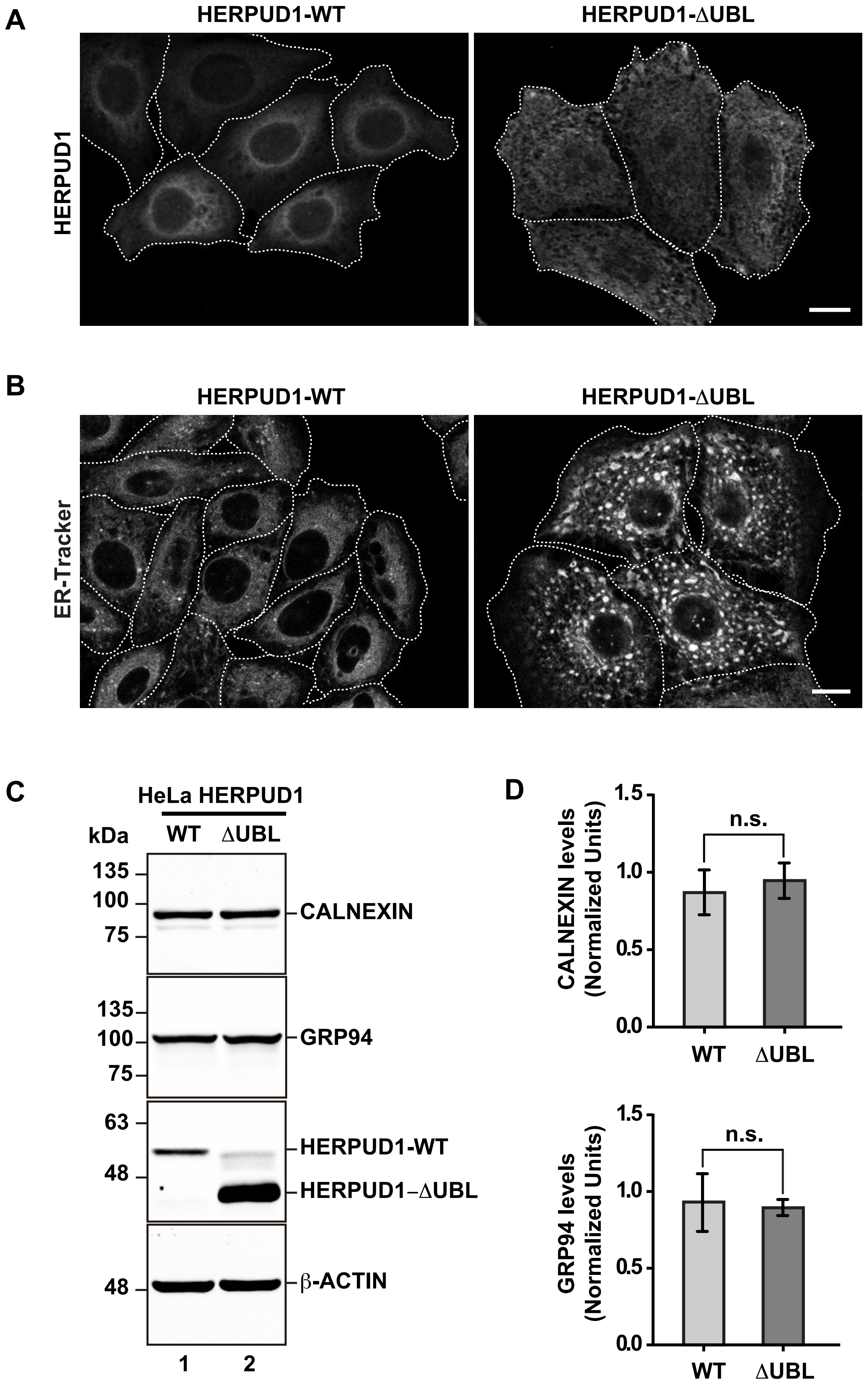


Supplementary Figure 3

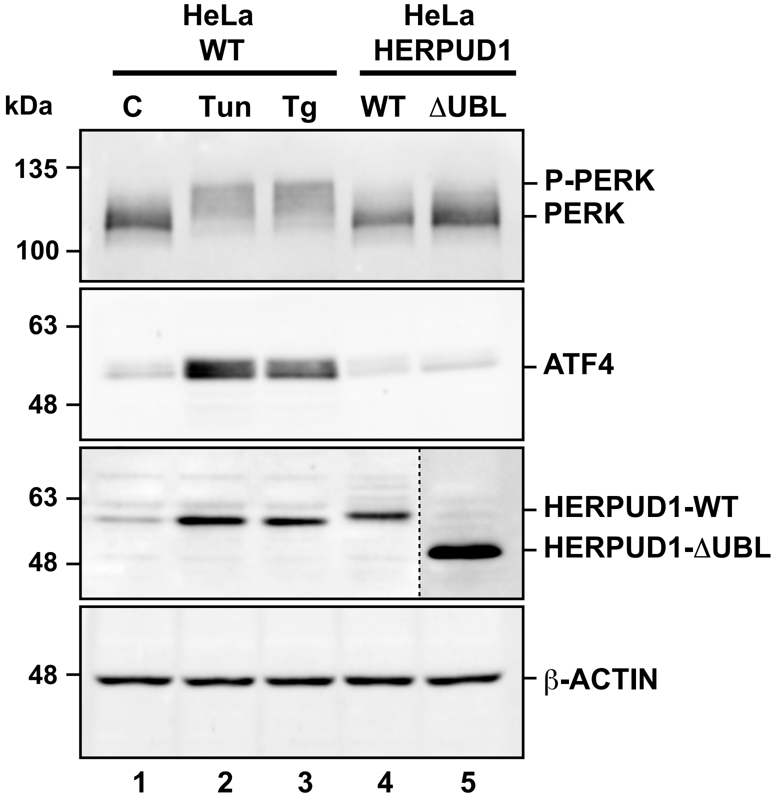


**Supplementary Figure 4**

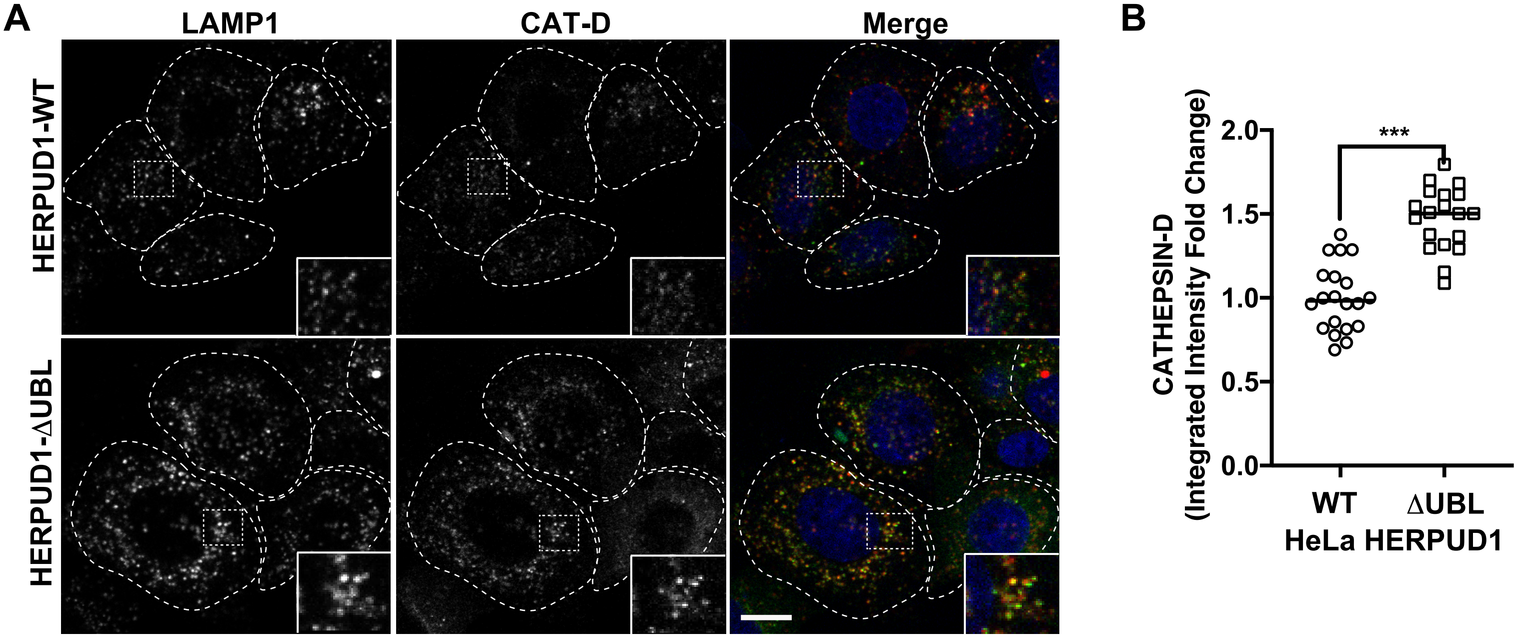


Supplementary Figure 5

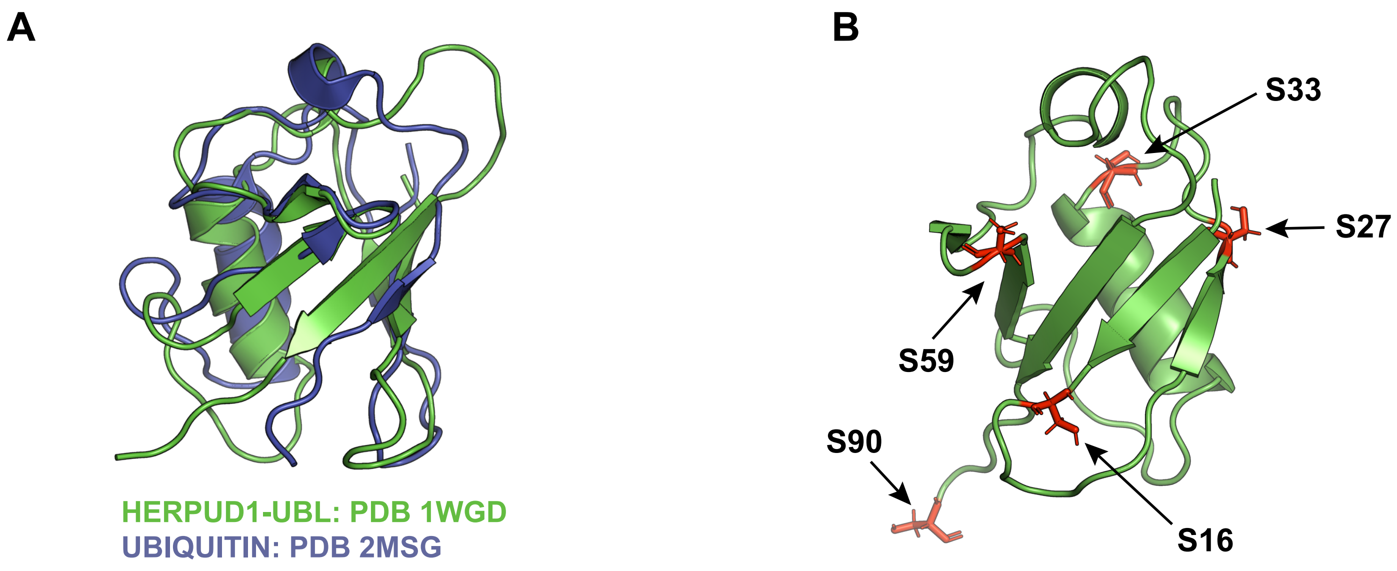


**Supplementary Figure 6**

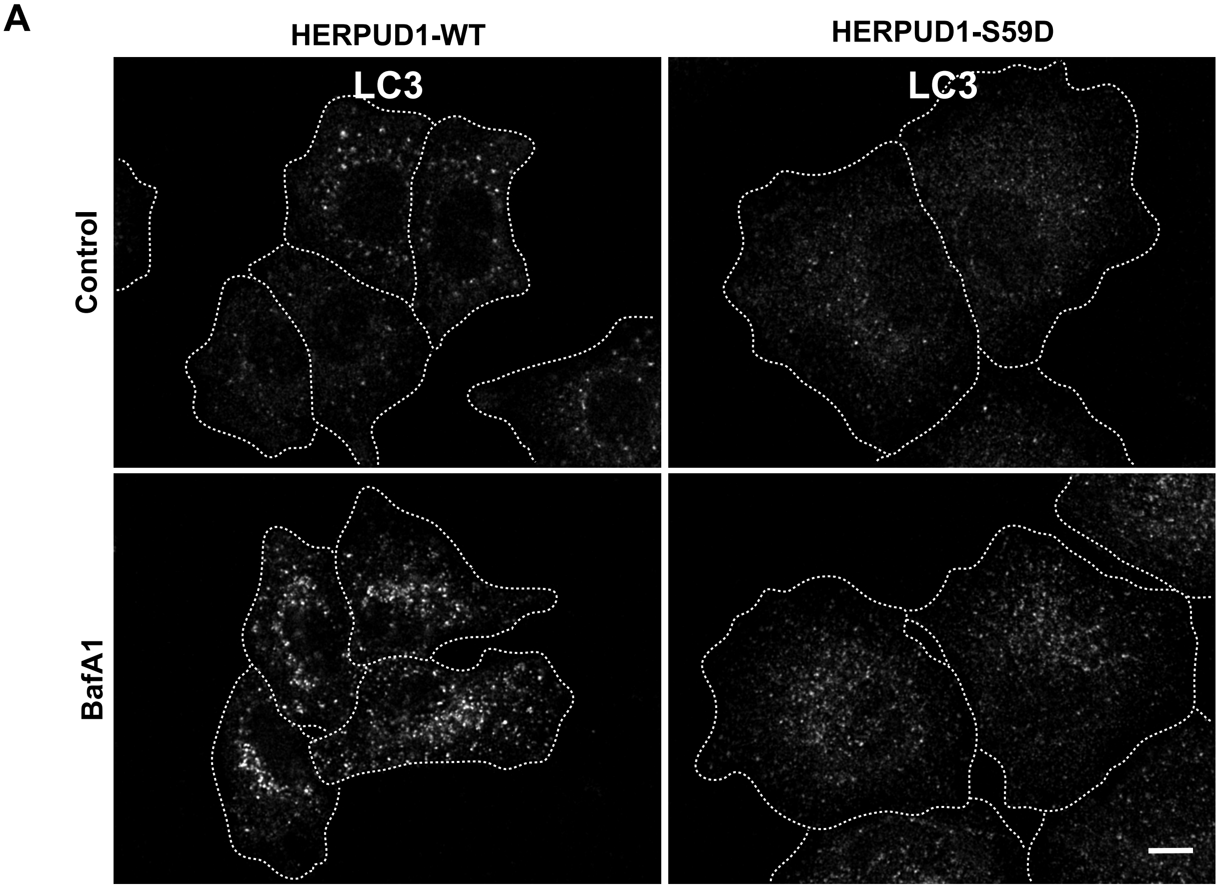


Supplementary Figure 7

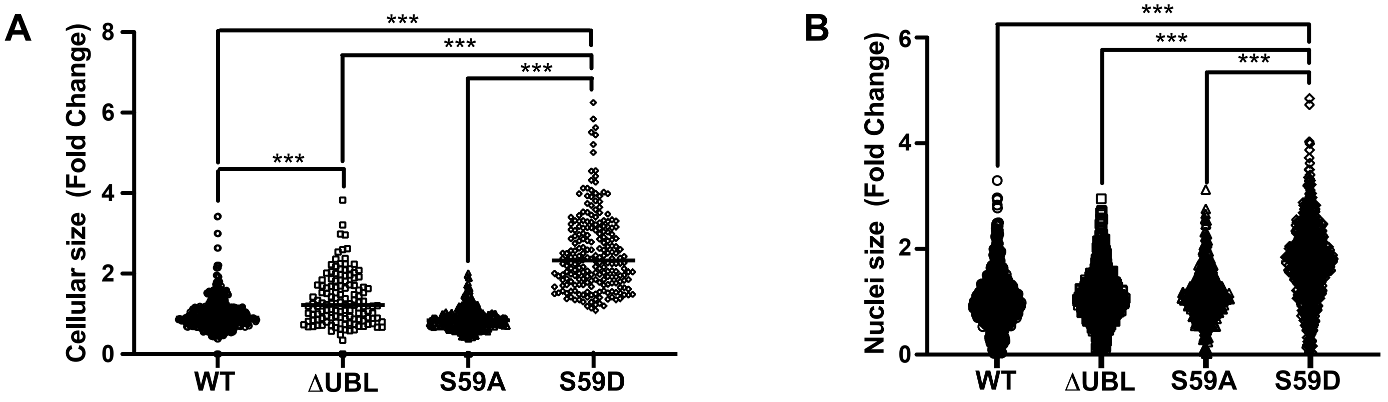


**Supplementary Figure 8**

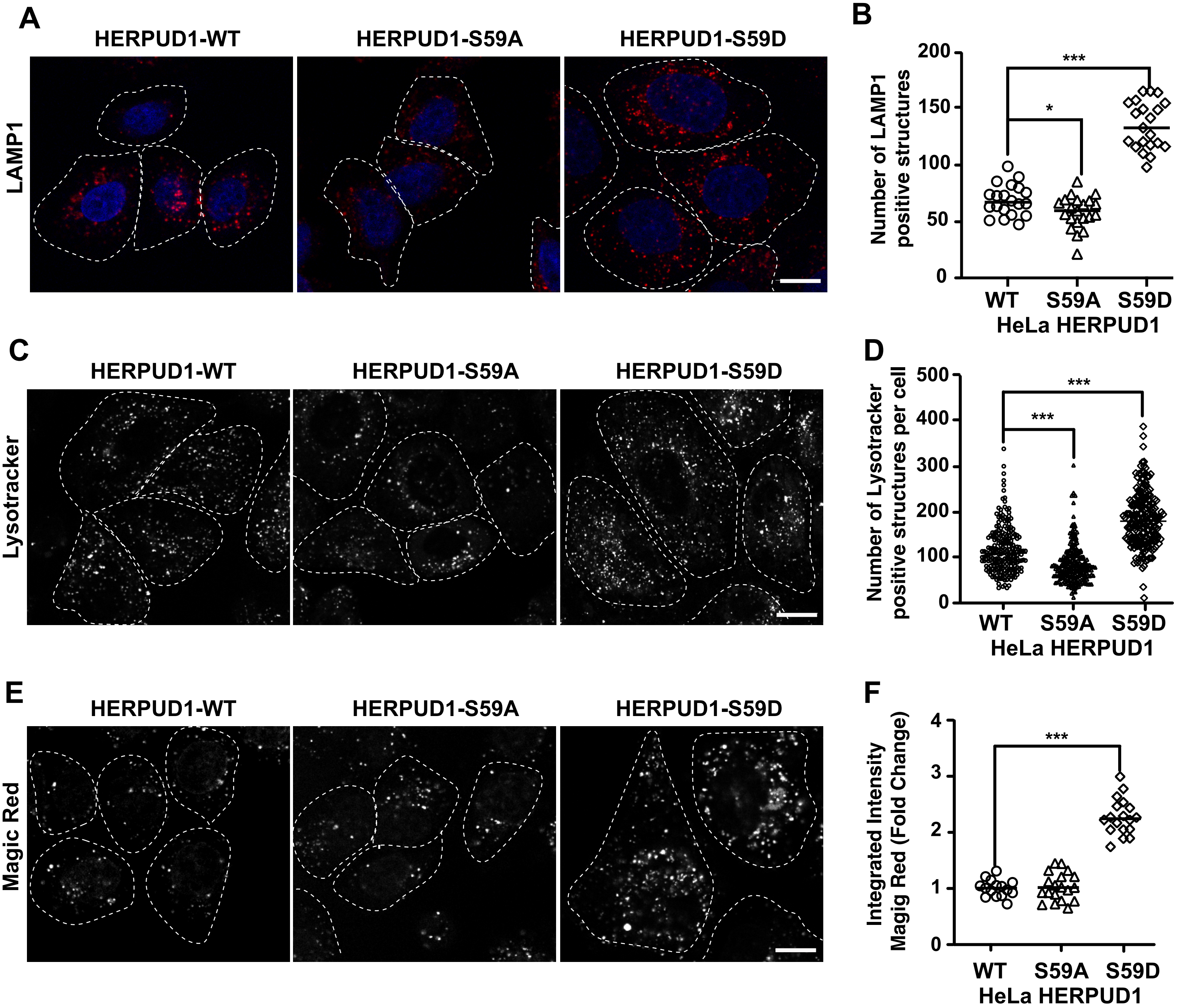


Supplementary Figure 9
